# Immunization with Herpes Simplex Virus Nanoparticles Targeting Both Attachment and Fusion Protect Against Infection

**DOI:** 10.64898/2026.04.24.720674

**Authors:** Dawid Maciorowski, Alexander C. Vostal, Wei Bu, Isabella S. Pytel, Sophia Antonioli-Schmit, Jianghai Zhu, Forrest H. Hoyt, Haotian Lei, Guangping Liu, Kristen Kaiser, Richard Herbert, Kennichi C. Dowdell, John T. Schiller, Kening Wang, Mark R. Howarth, Jeffrey I. Cohen

## Abstract

Herpes simplex virus 2 (HSV-2) is associated with genital ulcers, neonatal encephalitis, increased risk of HIV infection, and dementia. There is no licensed HSV-2 vaccine. We developed nanoparticles displaying the HSV-2 attachment protein gD and fusion mediation protein complex gH/gL. Immunization of mice and non-human primates elicited high levels of neutralizing antibodies. Vaccination conferred robust protection in mice, preventing disease and nearly eliminating infection and shedding following HSV-2 challenge. While gD induced high neutralizing antibody titers, gH/gL contributed substantially to protection despite lower neutralization titers. Instead, gH/gL immunization generated strong fusion-blocking responses which were an important correlate of protection, showing that standard neutralization assays incompletely capture the importance of fusion-blocking activity. These findings demonstrate that targeting both HSV-2 attachment and fusion elicit complementary mechanisms for protection from infection and that neutralizing antibody alone may be insufficient for protection. Overall, these results present an innovative strategy for an HSV-2 vaccine.

## INTRODUCTION

Prevention of herpes simplex virus type 2 (HSV-2) disease is a global health priority^1^, with an estimated 520 million people affected worldwide, with twice as many women infected as men^1,2^. During primary infection, HSV-2 infects epithelial cells in the genital tract, followed by infection of nerve endings and establishment of lifelong latency in the peripheral nervous system^3^. The virus frequently reactivates, resulting in recurrent painful genital lesions or asymptomatic viral shedding that facilitates transmission to sexual partners and neonates^3^. These episodes promote ongoing transmission, even in the absence of symptoms^4^. HSV-2 infection carries profound psychosocial consequences, increases the risk of HIV acquisition and transmission three- to four-fold, and globally causes ∼14,000 cases of neonatal herpes per year, though the incidence is likely substantially understated due to diagnostic and reporting limitations^5–8^. HSV-2 can cause severe disease in transplant recipients and patients with AIDS and has recently been implicated as a risk factor for Alzheimer’s disease^9–11^.

Despite long-term efforts, no HSV-2 vaccine has been licensed^12^. An effective HSV-2 vaccine has been considered a high priority by the United States (U.S.) National Institutes of Health and the U.S. Institute of Medicine^13,14^. Vaccine efficacy has been hindered by HSV’s robust immune evasion, its ability to directly infect the mucosa, and by the complex array of viral glycoproteins important for entry of the virus into cells^15,16^. HSV-2 entry into cells requires the coordinated action of four glycoproteins: gD, the gH/gL heterodimer, and gB^17^. Entry is initiated by the attachment of gD to host cell receptors, primarily nectin-1 and herpesvirus entry mediator (HVEM)^18,19^. Nectin-1 is broadly expressed, but enriched in epithelial cells and neurons, while HVEM is expressed in epithelial cells and enriched in lymphocytes^20,21^. Upon receptor engagement, gD interacts with gH/gL to initiate the membrane fusion cascade^22^. gH/gL is thought to act as a fusion mediator, interacting with gB to facilitate membrane fusion^23^. Among human herpesviruses, gD is unique to HSV, while gH/gL and gB are conserved among the other human herpesviruses^16^.

gD has been the most extensively targeted glycoprotein in HSV-2 vaccine clinical trials, yet no gD-based vaccine has achieved successful prevention of HSV-2 infection^12^. gD is a highly immunogenic glycoprotein and is the main target of neutralizing antibodies to HSV-2^24,25^ and several studies have demonstrated that gD-specific antibodies contribute to protection against HSV acquisition and disease^26–28^. Because of the critical roles of gD in receptor binding and initiation of fusion, as well as the ability of gD to induce neutralizing antibodies, gD likely remains an important component of an effective HSV vaccine^29–31^. Nonetheless, the largest clinical trial to date of a HSV-2 vaccine found that soluble gD was not effective at preventing HSV-2 disease and suggested that gD needs to be enhanced as an immunogen or combined with other viral glycoproteins to provide better protection^31^. This emphasis on gD has also reinforced serum neutralizing antibody titers as the principal benchmark for a prophylactic vaccine against HSV-2. However, HSV-2 entry is a multistep process requiring not only receptor engagement by gD but also downstream membrane fusion mediated by gH/gL and gB. Thus, antibody activities that interfere with fusion may not fully be represented by standard neutralization assays, and the extent to which such responses contribute to protection remains unresolved.

As an essential regulator of membrane fusion, gH/gL may elicit protective antibodies that operate through mechanisms distinct from those directed against gD. In contrast to gD, gH/gL has received comparatively little attention as an HSV-2 vaccine antigen. One early study demonstrated that immunization with soluble HSV-1 gH/gL elicited neutralizing antibodies, though to a lesser extent than soluble gD, and conferred protection comparable to gD in a dermal murine challenge model^32^. Another study evaluated whether soluble HSV-2 gH/gL and gB could enhance the protection from soluble HSV-2 gD in a guinea pig infection model, and found no benefit compared to gD alone^33^. Given the indispensable role of gH/gL in virus entry^16^ and the lack of effective HSV vaccines, we decided that further investigation of gH/gL as a vaccine immunogen was warranted, including whether protective immunity could be enhanced with nanoparticle assembly and whether protection is greater when gH/gL is co-administered with gD.

Multivalent nanoparticle display has emerged as a powerful strategy to enhance vaccine efficacy, promoting strong and long-lasting antibody responses by improving lymph node trafficking, B cell receptor cross-linking, and controlling antigen presentation for immunofocusing^34^. Virus-like particles (VLPs) and/or nanoparticles (NPs), including the licensed hepatitis B virus and human papillomavirus vaccine have good safety profiles, induce durable immunity, and have had substantial clinical success^35^. We employed a plug-and-display methodology to overcome existing challenges of NP display of HSV antigens by harnessing efficient and irreversible isopeptide bond formation between SpyTag003^36^ fused to HSV-2 antigens and SpyCatcher003 (SC3) fused to a computationally designed mi3 60-mer nanocage^37^.

Here, we show that NP display of HSV-2 antigens provides enhanced protection compared to their corresponding soluble forms. Although gD NPs elicited substantially higher neutralizing antibody titers than gH/gL NPs, gH/gL NPs induced stronger fusion-blocking antibody responses and resulted in lower HSV-2 shedding after HSV-2 challenge. Immunization of NPs displaying gD and gH/gL led to high titer neutralizing and fusion-blocking antibody responses in both mice and non-human primates. Co-administering NPs displaying gD and gH/gL followed by a lethal HSV-2 challenge eliminated disease and prevented infection and shedding in most mice. This dual NP vaccine candidate demonstrates that simultaneously targeting both the viral attachment and fusion apparatus leads to enhanced protective responses and provides a promising pathway towards an effective HSV-2 vaccine.

## Results

### Construction and Characterization of gD- and gH/gL-Nanoparticles

High molecular weight antigens bearing multiple post-translational modifications have often been challenging to assemble onto nanoparticles (NPs), resulting in aggregation, heterogeneous particles, or instability^38^. Our previous attempts at genetic fusion of HSV-2 gH/gL to *Helicobacter pylori*-ferritin^39^ for nanoparticle assembly were unsuccessful, likely due to the steric hindrance of gH/gL on the ferritin monomers self-assembling into nanoparticles (data not shown). We hypothesized that allowing SC003-mi3 NPs to assemble, before conjugating the bulky gH/gL antigen to the particles using the SpyTag/SpyCatcher system, would result in successful production of gH/gL NPs (**Figure 1A**). NPs with SpyCatcher (SC003-mi3 without a conjugated glycoprotein) were expressed in *Escherichia coli* (**Supplemental Figure 1A, 1B**). HSV-2 gD and gH constructs retained the endogenous signal sequence and their ectodomains were fused to a flexible Gly/Ser linker and SpyTag003 at the C-terminus (**Figure 1B).** gHSpyTag003 was co-transfected with full-length HSV-2 gL and expressed in mammalian Expi293F cells (**Figure 1B**).

**Figure 1.**
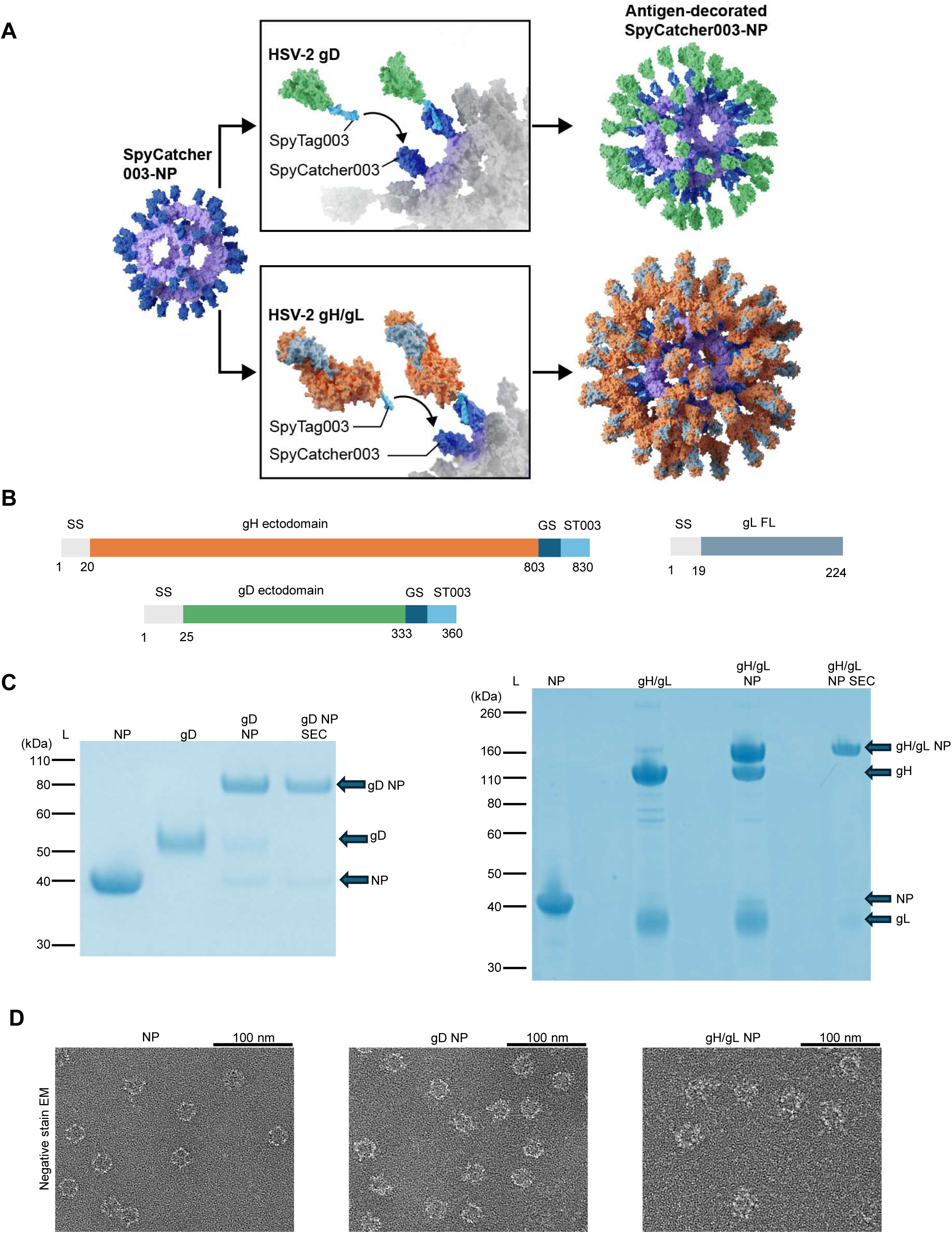
Construction and Characterization of Nanoparticles (NPs). A) Schematic of HSV-2 glycoproteins coupling to NPs. HSV-2 glycoprotein-SpyTag fusion constructs (gD based on PDB 4MYV or gH/gL based on PDB 3M1C) are coupled to SpyCatcher003 mi3 NPs. B) Diagram of HSV-2 gD, gH, and gL constructs. gD and gH ectodomains are fused at their carboxyl termini to SpyTag003 (ST003). A plasmid expressing gL full-length (FL) was co-transfected with the gH plasmid at a 1:1 molar ratio to produce gH/gL. GS is a glycine-serine linker. C) SDS-PAGE with Coomassie staining of unconjugated NPs, soluble gD and gH/gL, NPs conjugated to gD or gH/gL before or after purification by size exclusion chromatography (SEC). L is size marker ladder. Gels converted to Coomassie blue color in PowerPoint. D) Negative stain electron microscopy images of NPs, gD NPs, and gH/gL NPs.

SpyTag-fusion proteins were purified from the supernatant via SpySwitch affinity chromatography^40^ (**Supplemental Figure 1C, 1D**). We coupled gD to SC003-mi3 at a 1.5:1 molar ratio, with high efficiency of covalent coupling shown by SDS-PAGE (**Figure 1C, left panel, Supplemental Figure 2A left panel**). We conjugated HSV-2 gH/gL to NPs to near saturation, using a 3:1 molar ratio of gH/gL-SpyTag003:SC003-mi3 **(Figure 1C, right panel, Supplemental Figure 2A, right panel**). NP, gD NP, and gH/gL NP were then purified by size-exclusion chromatography (SEC) (**Supplemental Figure 3A-C**). NP, gD NP, and gH/gL NP were visualized with negative stain transmission electron microscopy and showed well-formed particles (**Figure 1D, Supplemental Figure 1E**). Cryogenic electron microscopy (CryoEM) was used to attempt to create a 3D reconstruction of gD NP and gH/gL NP. CryoEM resolved the gH/gL NP core to a 3.42 Å resolution with protrusions localized to where gH/gL is conjugated. Defined density, corresponding to the conjugated gD (data not shown) or gH/gL glycoproteins, was not resolved **(Supplemental Figure 4A-F**), even after masking procedures, likely due to the flexibility of the linkers. Nanoparticle hydrodynamic radius (HDR) and homogeneity were measured with dynamic light scattering. Uncoupled NP had a HDR of 17.6 ± 1.9 nm, gD NPs had a HDR of 19.0 ± 1.4 nm, and gH/gL NPs had a HDR of 27.0 ± 1.8 nm (**Supplemental Figure 3D**). These data indicate that the NPs were predominantly homogeneous single particles, and had the expected radius shifts after antigen coupling^41^.

### gD and gH/gL Nanoparticle Vaccines Induce Potent Neutralizing and Fusion-Blocking Antibody Responses in Mice, Despite Comparable Binding Antibody Titers

To evaluate immunogenicity of nanoparticle (NP) and soluble vaccines, BALB/c mice were immunized intramuscularly (IM) with phosphate-buffered saline (PBS) or 5 µg of soluble gD, soluble gH/gL, soluble gD+gH/gL, gD NP, gH/gL NP, or gD NP+gH/gL NP (**Figure 2A**). gD and gH/gL were given at an equimolar ratio when used in combination. Each vaccine as well as the PBS group was combined with Sigma Adjuvant Systems (SAS) adjuvant^42^ at the time of immunization. A homologous boost was given 3 weeks later. Serum-antibody titers to gD and gH/gL were measured using enzyme-linked immunosorbent assays (ELISA) 1.5 weeks post dose 2. Both anti-gD and anti-gH/gL ELISA antibody titers to the corresponding soluble antigen were consistently high and similar to gD NP and gH/gL NP titers among the vaccinated groups, with no comparisons reaching statistical significance (**Figure 2B, 2C**). Importantly, combining gD and gH/gL caused no loss of antibody response to either gD or gH/gL. We next measured HSV-2 serum neutralizing titers (dilution of serum that inhibits infection by 50% [IC_50_]) in Vero cells. Of the soluble glycoprotein vaccines, soluble gH/gL had the lowest mean neutralizing antibody titers, ∼3-fold lower compared to soluble gD (47 vs. 133, respectively, p<0.0001) and ∼2-fold lower compared to soluble gD+gH/gL (47 vs. 103, respectively, p=0.0027) (**Figure 2D**). With all glycoproteins, NPs elicited higher neutralizing antibody titers compared to their corresponding soluble glycoprotein counterparts. The virus neutralization titer was ∼31-fold higher in mice immunized with gD NPs than those immunized with soluble gD (4,119 vs. 133, respectively, p<0.0001) and mice receiving gH/gL NPs elicited ∼11-fold higher titers than with soluble gH/gL (524 vs. 47, respectively, p<0.0001). Immunization with gD NP+gH/gL NP elicited ∼23-fold higher titers than the soluble combination (2,397 vs. 103, respectively, p<0.0001). Among groups receiving NP vaccines, neutralizing antibody titers were highest in the gD NP-vaccinated group, ∼8-fold higher titers than gH/gL NP (4,119 vs. 524, respectively, p<0.0001) and ∼2-fold higher titers than gD NP+gH/gL NP (4,119 vs. 2,397, respectively, p=0.002). The results from the neutralization assay showed that while NP display of gD and gH/gL increases the neutralizing antibody titers compared to their soluble forms, gD is a better target than gH/gL for inducing neutralizing antibodies via the standard Vero cell-based neutralization assay.

**Figure 2.**
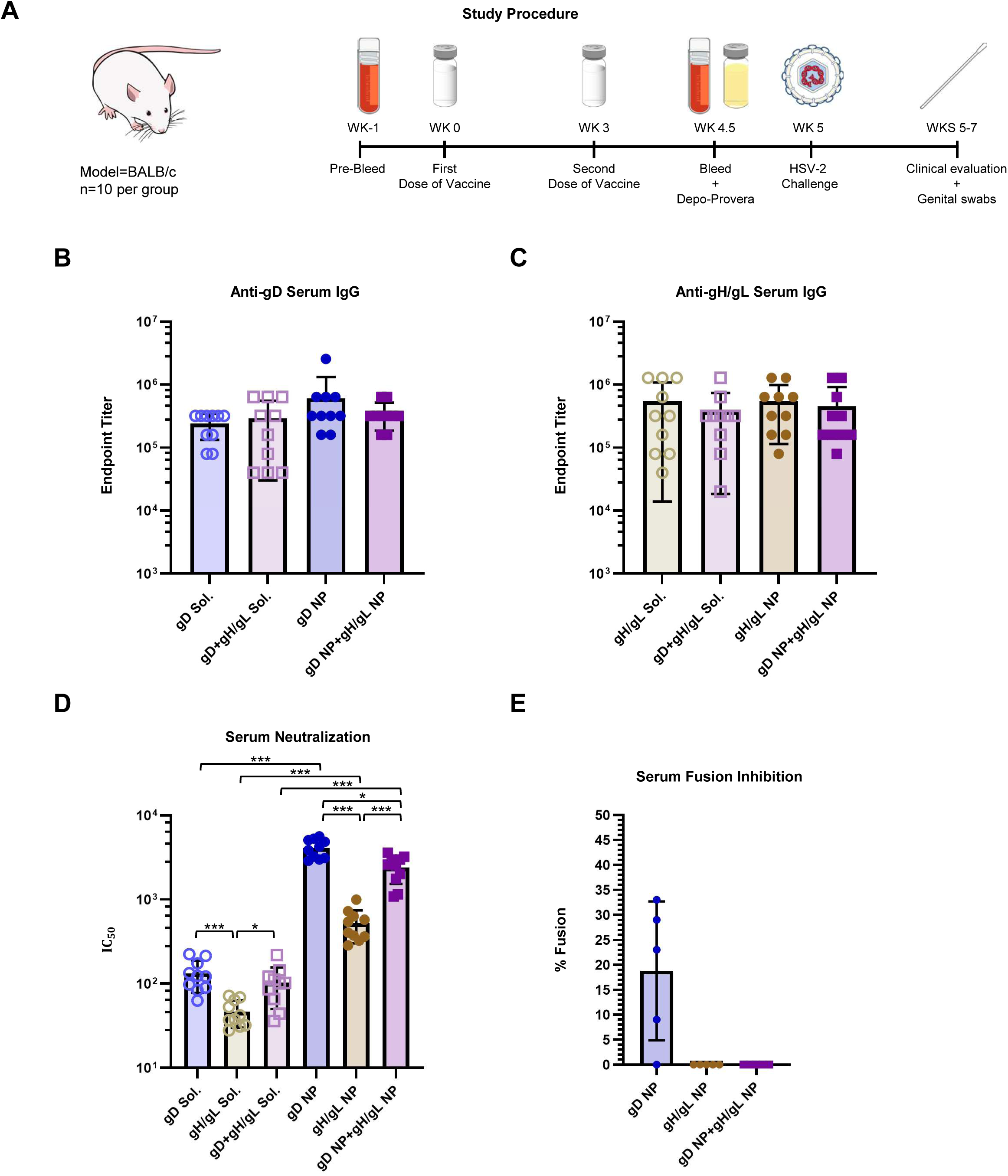
Antibody responses in mice Immunized with soluble gD, gH/gL, or gD+gH/gL, or with gD NP, gH/gL NP, or gD NP+gH/gL NP. A) Schematic of the mouse vaccination study. BALB/c mice (n=10 per group) were immunized intramuscularly with 2 doses of vaccines at 5 µg per dose in SAS adjuvant given 3 weeks apart. Serum was analyzed from mice bled 1.5 to 2 weeks after the second dose of vaccine. B) HSV-2 gD binding ELISA antibody titers in serum shown as endpoint titers. Bars indicate the means and error bars indicate standard deviations. None of the differences between the titers were significant. C) HSV-2 gH/gL binding ELISA antibody titers in serum shown as endpoint titers Bars indicate the means and error bars indicate standard deviations. None of the differences between the titers were significant. D) HSV-2 neutralizing antibody titers in serum. IC_50_ is the dilution of sera that inhibits infection by 50 Bars indicate the means and error bars indicate standard deviations. *p <0.0056; ***p<0.0001. E) HSV-2 fusion inhibition by antibodies in serum (n=5 per group). Bars indicate the means and error bars indicate standard deviations. Sera from mice before immunization were used as a positive control, while using Chinese hamster ovary (CHO)-K1 cells expressing HSV-2 gH/gL and luciferase without gB were used as a negative control for fusion. Percent fusion was defined as the luminescence of immune serum samples divided by the luminescence of pooled pre-immune serum samples x 100. After correcting for multiple comparisons (0.05/2 comparisons=0.025), the differences between gD NP vs. gH/gL NP and gD NP vs. gH/gL NP trended towards significance (p=0.038).

We next investigated whether sera from mice immunized with NPs have fusion-blocking activity and hypothesized that if the main role of gH/gL is in virus fusion, then sera from mice immunized with gH/gL NP would exhibit better fusion-blocking than mice receiving gD NP. While sera from mice immunized with gD NP inhibited HSV-2 fusion by ∼82%, sera from mice immunized with either gH/gL NP or gD NP+gH/gL NP completely inhibited fusion (**Figure 2E**). Thus, these data suggest that immunization with gD and gH/gL allows for complementary immune targeting of both virus attachment and virus fusion.

### Targeting Both HSV-2 Attachment and Fusion Nearly Eliminates HSV-2 Vaginal Shedding

As a stringent test of efficacy of gD and gH/gL NP vaccinations in protecting against intra-vaginal HSV-2 challenge, mice were inoculated with 64,000 plaque forming units (PFU) of HSV-2 strain 333, which is approximately 63 times the 50% lethal dose (LD_50_). All mice that received PBS control died by day 10 (disease score of 6), while none of the mice that received vaccine candidates died or developed systemic disease (**Figure 3A**). Starting at day 5 post-challenge, mice were followed daily for clinical signs, such as vaginal erythema, genital lesions, genital hair loss, ruffled fur, abnormal gait, hunched back, hind-limb paralysis, or lethargy. By day 7, all PBS control mice developed lesions with marked vaginal erythema. In mice receiving soluble gD, one mouse developed genital lesions, and two mice developed vaginal erythema. In the soluble gH/gL group, one mouse developed genital lesions, and three mice had vaginal erythema. Three mice receiving the soluble gD+gH/gL combination had vaginal erythema. One mouse immunized with gD NPs developed vaginal erythema while mice receiving gH/gL NPs and gD NP+gH/gL NP did not develop any clinical disease (**Figure 3A**).

**Figure 3.**
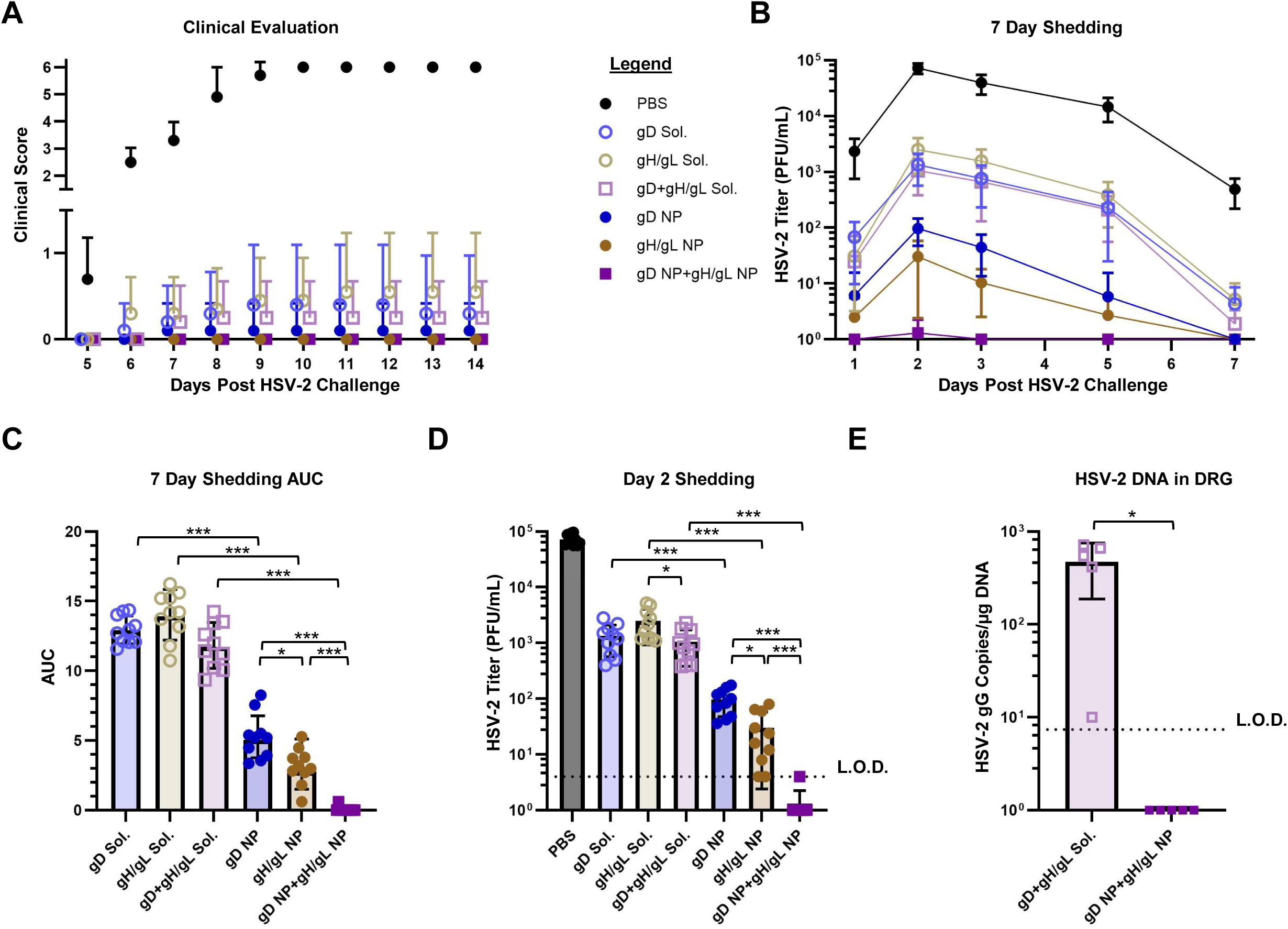
Protection of mice immunized with soluble gD, gH/gL, gD+gH/gL, or with gD NP, or gH/gL NP, or gD NP+gH/gL NP from HSV-2 vaginal challenge. BALB/c mice (n=10 per group) were immunized intramuscularly with 2 doses of vaccine (5 µg) in SAS adjuvant, 3 weeks apart. 2 weeks later, mice were intravaginally challenged with 64,000 PFU of HSV-2 strain 333. Data presented are from three independent experiments. (A) Mice were monitored daily for clinical signs of HSV-2 infection including vaginal erythema, genital lesions, genital hair loss, ruffled fur, lethargy, abnormal gait, hunched back, or hind-limb paralysis and given a score of 0 (no disease) to 6 (dead) (see Methods). Data points indicate the means and error bars indicate standard deviations. There were no statistical differences in groups receiving soluble glycoproteins or NPs. (B) HSV-2 titers from vaginal swabs taken on days 1, 2, 3, 5, and 7 post-challenge. Data points indicate the means and error bars indicate standard deviations. (C) Plotted AUC of the HSV-2 titers from vaginal swabs over the 7 day time course in panel B. Bars indicate the means and error bars indicate standard deviations. *p <0.0056; *** p<0.0001. (D) HSV-2 titers from vaginal swabs of individual mice on day 2. Bars indicate the means and error bars indicate standard deviations. L.O.D. is the limit of detection defined as 4 PFU/mL. *p <0.0056; *** p<0.0001. (E) HSV-2 DNA copy number in sacral dorsal root ganglia (DRG) of mice obtained 14 days after intravaginal challenge with HSV-2 (n=5 mice per group). Bars indicate the means and error bars indicate standard deviations. Copy numbers were determined by qPCR using an HSV-2 gG probe. L.O.D. is the limit of detection defined as 7.4 copies of HSV-2 DNA. *p <0.05.

Virus shedding post-intravaginal challenge represents a more stringent test of vaccine efficacy than disease score or mortality, providing an important metric for whether new immunogens may outperform vaccines that were not successful in clinical trials^43^. On days 1, 2, 3, 5, and 7 post-challenge, vaginal swabs were obtained from mice and evaluated for infectious HSV-2. Mean vaginal titers and area under the curve (AUC) were plotted for each group over the 7-day evaluation (**Figure 3B, 3C**). All mice immunized with NPs had lower virus shedding than their soluble glycoprotein forms (soluble gD vs. gD NP, p<0.0001, soluble gH/gL vs. gH/gL NP, p<0.0001, soluble gD+gH/gL vs. gD NP+gH/gL NP, p<0.0001). Importantly, mice vaccinated with gH/gL NP had lower shedding than mice receiving gD NP (p=0.0051) even though gH/gL NP elicited lower neutralizing antibody titers than gD NP (**Figure 2D, 3B, 3C).** Although higher neutralizing antibody titers correlated with protection when comparing soluble antigens to their NP counterparts, protection between gD NP and gH/gL NP was more correlative with fusion-blocking antibodies (**Figure 2D, 2E, 3B, 3C**).The gD NP+gH/gL NP combination had strong neutralizing antibody responses (**Figure 2D**) and potent fusion-blocking activity (**Figure 2E**) and outperformed both gD NP and gH/gL NP alone, as immunization with the combination vaccine nearly eliminated shedding (gD NP+gH/gL NP vs. gD NP, p<0.0001, gD NP+gH/gL NP vs. gH/gL NP, p<0.0001).

Virus shedding in individual mice was plotted on day 2 and 5 post-challenge (**Figure 3D, Supplemental Figure 2B**). On day 2, the virus titers reached the peak over the time course with mice receiving the PBS control having a mean titer of 72,500 PFU/mL. Among the groups receiving soluble vaccines, soluble gD+gH/gL had lower virus titers than soluble gH/gL (1,050 vs. 2,490, respectively, p=0.0060). All groups receiving NPs had lower virus titers than groups receiving the soluble glycoprotein form (gD NP vs. soluble gD, 96.8 vs. 1,340, respectively, p<0.0001; gH/gL NP vs. soluble gH/gL, 30.2 vs. 2,490, respectively, p<0.0001; gD NP+gH/gL NP vs. soluble gD+gH/gL, 0.4 vs. 1,050, respectively, p<0.0001). Differences in HSV-2 titers between gD NP and gH/gL NP were significant (96.8 vs. 30.2, respectively, p=0.0019). In the group that received co-administered gD NP+gH/gL NP, only one of the ten mice had detectable infectious HSV-2 over the entire 7 day time course, where on day 2 post challenge, one mouse had a virus titer just reaching the assays limit of detection of 4 PFU/mL (**Figure 3B, 3C, 3D**). gD NP+gH/gL NP showed superior protection by nearly eliminating infection, while all mice in the gD NP and gH/gL NP groups had detectable virus, albeit low titers (gD NP+gH/gL NP vs. gD NP, 0.4 vs. 96.8, respectively, p<0.0001; gD NP+gH/gL NP vs. gH/gL NP, 0.4 vs. 30.2, respectively, p<0.0001). On day 5 post-challenge, mice immunized with soluble glycoproteins, gD, gH/gL, and gD+gH/gL had detectable virus with mean titers ∼282 PFU/mL (230.4, 379.6, 210.2 PFU/mL, respectively) (**Supplemental Figure 2B**). Detectable virus shedding was present in 4 of 10 mice receiving gD NP, 3 of 10 receiving gH/gL NP, and none receiving gD NP+gH/gL NP. Each NP group had significantly lower titers when compared to their soluble glycoprotein counterparts (p<0.0001, p<0.0001, p<0.0001 for gD, gH/gL, and gD+gH/gL, respectively).

HSV-2 establishes latency in sacral dorsal root ganglia (DRG) after primary infection, from which HSV-2 can subsequently reactivate. To determine whether our combined gD NP+gH/gL NP vaccine could prevent HSV-2 from infecting DRG, we used quantitative polymerase chain reaction (qPCR) to measure the level of HSV-2 DNA in DRG after challenge. DRGs were harvested from mice vaccinated with soluble gD+gH/gL or gD NP+gH/gL NP on day 14 post-challenge. No sacral DRG from mice vaccinated with gD NP+gH/gL NP had detectable HSV-2 DNA, while 5 of 5 mice vaccinated with soluble gD+gH/gL had HSV-2 DNA in their sacral DRG (**Figure 3E)**. Mice receiving PBS control were not tested, since all had died before day 14.

Taken together, these results indicate that vaccination with NPs displaying gD and gH/gL afforded better protection against HSV-2 challenge compared to the corresponding soluble glycoproteins. After HSV-2 challenge, mice receiving gH/gL NP had lower virus titers in the vaginal tract than gD NP, even though gH/gL NP elicited substantially lower neutralizing titers on a standard Vero cell-based assay. The discordance between neutralizing antibody titers and in vivo protection suggests that standard neutralization assays do not fully capture the spectrum of protective antibody functions elicited by vaccination, specifically fusion-blocking antibodies. Overall, immunization with HSV-2 glycoprotein NPs targeting both the viral attachment and fusion apparatus resulted in superior protection, nearly eliminating vaginal shedding and preventing HSV-2 entry into DRG.

### Co-administered gD NPs and gH/gL NPs are more Protective than Either gD NP or gH/gL NP Alone

With the near complete protection from infection granted by gD NP+gH/gL NP, but not either gD NP or gH/gL NP alone, we hypothesized that optimal protection required equal presentation of both gD and gH/gL glycoproteins, and that diluting one component would decrease protection. To evaluate this, mice were immunized following the same protocol as before (**Supplemental Figure 5A**) with PBS, or 5 µg of total antigen per dose of gD NP, gH/gL NP, or a combination of gD NP+gH/gL NP at 3 different molar ratios of gD NP to gH/gL NP (10:1, 3:1, and 1:1, respectively). 1.5 weeks post dose 2, mice were bled and serum neutralizing antibody titers were evaluated (**Supplemental Figure 5B**). As expected, gD NP elicited higher neutralizing antibody titers than gH/gL NP (4243 vs. 468, respectively, p<0.0001) and the groups that included both gD NP+gH/gL NP at 10:1, 3:1, 1:1 molar ratios, had higher neutralizing antibody titers than gH/gL NP alone (3556 vs. 468, respectively, p<0.0001; 2252 vs. 468, respectively, p<0.0015; 2626 vs. 468, respectively, p=0.0001).

Two weeks post dose 2, mice were challenged with HSV-2 and by day 9, mice that received the PBS control died (**Supplemental Figure 5C**). Due to the efficacy of experimental vaccine groups, clinical scoring was not stringent enough of an evaluation to observe differences between groups. Mice were swabbed on days 1, 2, 3, 5, and 7 post-challenge to evaluate virus shedding in the vaginal tract, and both titers and AUC were plotted (**Supplemental Figure 5D, 5E**). As the combination of gD NP+gH/gL NP was closer to equimolar presentation, the difference in shedding became statistically significant (gD NP vs. 3:1 molar ratio, p=0.0033; gD NP vs. 1:1 molar ratio, 0.0032).

In summary, these findings indicate that co-administration of gD NPs and gH/gL NPs is important to obtain protection from HSV-2 infection and that the two vaccines provide at least an additive protective benefit compared to immunization with either NP alone.

### HSV-2 NPs Induce Durable Humoral Responses that Protect from Disease

Next, we evaluated the duration of protective immune responses afforded by gD NP+gH/gL NP immunization. Mice received the same regimen and dosage as in the previous experiments, but only the equimolar combination vaccines were evaluated (**Figure 4A, 4B**). After the second dose, mice were bled at different points over 8 months. To evaluate the time-course of the antibody response, serum neutralization and glycoprotein binding were determined. Over the 8-month time course, mice immunized with gD NP+gH/gL NP maintained higher levels of neutralizing antibodies than mice receiving soluble gD+gH/gL (**Figure 4C**). One-month post-dose 2, mice immunized with gD NP+gH/gL NP elicited ∼17-fold higher neutralizing antibody titers than soluble gD+gH/gL (2,284.2 vs. 130.8, respectively, p=0.0079). These comparatively high neutralizing antibody titers generated by gD NP+gH/gL NP were better maintained throughout the study, since at 8 months post-dose 2 only 2 mice immunized with soluble gD+gH/gL had detectable neutralizing antibodies (551 vs. 5.6, respectively, p=0.0079). While serum anti-gD binding antibody levels were similar between gD NP+gH/gL NP and soluble gD+gH/gL at one-month post-dose 2, by 8 months post-dose 2 the gD NP+gH/gL NP group had ∼13-fold higher anti-gD serum titers than the soluble gD+gH/gL group (44,800 vs 3,400, respectively, p=0.0079) (**Figure 4D**). Over the time course, mice immunized with soluble gD+gH/gL had an observed ∼18-fold reduction in anti-gD IgG titers, while mice immunized with gD NP+gH/gL NP had a ∼4-fold reduction in anti-gD titers.

**Figure 4.**
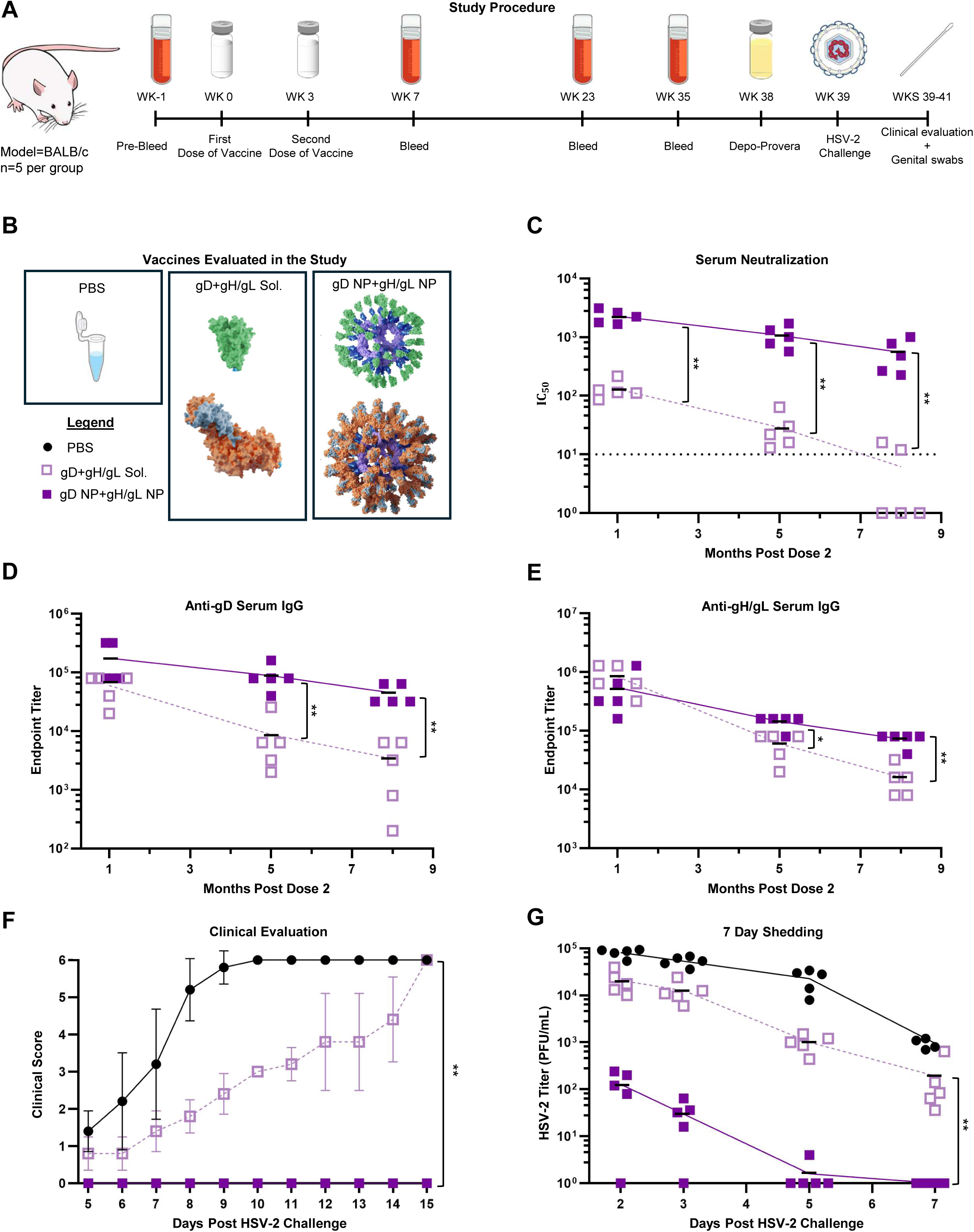
Protection of mice immunized with soluble gD+gH/gL, or gD NP+gH/gL NP from HSV-2 vaginal challenge and durability of antibody responses. BALB/c mice (n=5 per group) were immunized IM with 2 doses of vaccines (5 µg) in SAS adjuvant given 3 weeks apart. Mice sera were collected at various intervals. 9 months post-dose 2, mice were challenged intravaginally with 64,000 PFU of HSV-2 strain 333. (A) Study procedure of the long-term challenge. (B) Schematic of vaccines evaluated in the study. (C) Neutralizing antibody titers in sera from mice at months 1, 5, and 8 post vaccine dose 2. IC_50_ is the dilution of sera that inhibits infection by 50%. Data points indicate individual mice and bars indicate means. **p<0.01. (D) HSV-2 gD ELISA antibody titers in sera collected from mice at months 1, 5, and 8 post vaccine dose 2. Data points indicate individual mice and bars indicate means. **p<0.01. (E) HSV-2 gH/gL ELISA antibody titers in sera collected from mice at months 1, 5, and 8 post vaccine dose 2. Data points indicate individual mice and bars indicate means. *p<0.05, **p<0.01. (F) Mice were monitored daily for clinical signs of HSV-2 infection including vaginal erythema, genital lesions, genital hair loss, ruffled fur, lethargy, abnormal gait, hunched back, or hind-limb paralysis and given a score of 0 (no disease) to 6 (dead) (see Methods). Data points indicate individual mice and bars indicate means. **p<0.01. (G) HSV-2 titers from vaginal swabs taken on days 2, 3, 5, and 7 post-challenge with HSV-2. Data points indicate individual mice and bars indicate means. **P□<□0.01.

Like anti-gD antibody responses, anti-gH/gL serum binding antibody responses between soluble gD+gH/gL and gD NP+gH/gL NP were similar when measured 1 month post dose 2 (**Figure 4E**). At 5 months post dose 2, titers waned more significantly in mice receiving soluble gD+gH/gL than gD NP+gH/gL NP (60,000 vs. 144,000, respectively, p=0.032) and the difference became greatest at 8 months post dose 2 (16,000 vs. 72,000, respectively, p=0.0079). Over the time course, mice immunized with soluble gD+gH/gL had a ∼52-fold reduction in anti-gH/gL IgG titer, while mice immunized with gD NP+gH/gL NP had a ∼8-fold reduction in titers.

9 months after dose 2, mice were challenged and evaluated as described previously. All PBS control mice died by day 10 and all mice immunized with soluble gD+gH/gL died by day 15 (**Figure 4F**). No mouse receiving gD NP+gH/gL NP died or developed any signs of disease. Vaginal swabs were evaluated for infectious virus on days 2, 3, 5, and 7 post challenge (**Figure 4G**). On days 2 and 3, for mice receiving gD NP+gH/gL NP, 4 out of 5 mice had detectable infectious virus; the mean virus titer was ∼163-fold lower on day 2 for the nanoparticle group than the soluble glycoprotein group (128 vs. 20,800, respectively). By day 5, 1 mouse out of 5 immunized with gD NP+gH/gL NP had detectable virus and by day 7 all these mice were negative for infectious virus. Compared to gD NP+gH/gL NP, all mice receiving the soluble gD+gH/gL had high viral titers for the 7-day time course (p=0.0079).

Overall, 9 months after dose 2, mice immunized with gD NP+gH/gL NP and challenged with HSV-2 were protected from death and disease, displayed marked reduction in vaginal shedding, and by day 5 shedding was undetectable in all mice except one. These data show that NP display of HSV-2 glycoproteins affords more durable protection than their soluble glycoprotein forms.

### Passive Transfer of IgG from gD NP+gH/gL NP Vaccinated Mice Protects Naive Mice Against HSV-2 Challenge

To evaluate the protective efficacy of immune sera elicited by gD NP+gH/gL NP, BALB/c mice were passively immunized and subsequently challenged with HSV-2 (**Figure 5A**). To obtain purified IgG, mice were immunized IM 2 times, 3 weeks apart with PBS, 5 µg of soluble gD+gH/gL, or gD NP+gH/gL NP. Mice were then terminally bled 2 weeks post-dose 2; serum was collected and pooled and underwent Protein G purification. Mice received IgG (40 µg total IgG per gram of body weight) intraperitoneally (IP) on days -3, -1, 0 of challenge. Following the final IP administration, mice were challenged intravaginally with 64,000 PFU (63 x LD_50_) of HSV-2 strain 333 and then weighed and evaluated for clinical disease and shedding.

**Figure 5.**
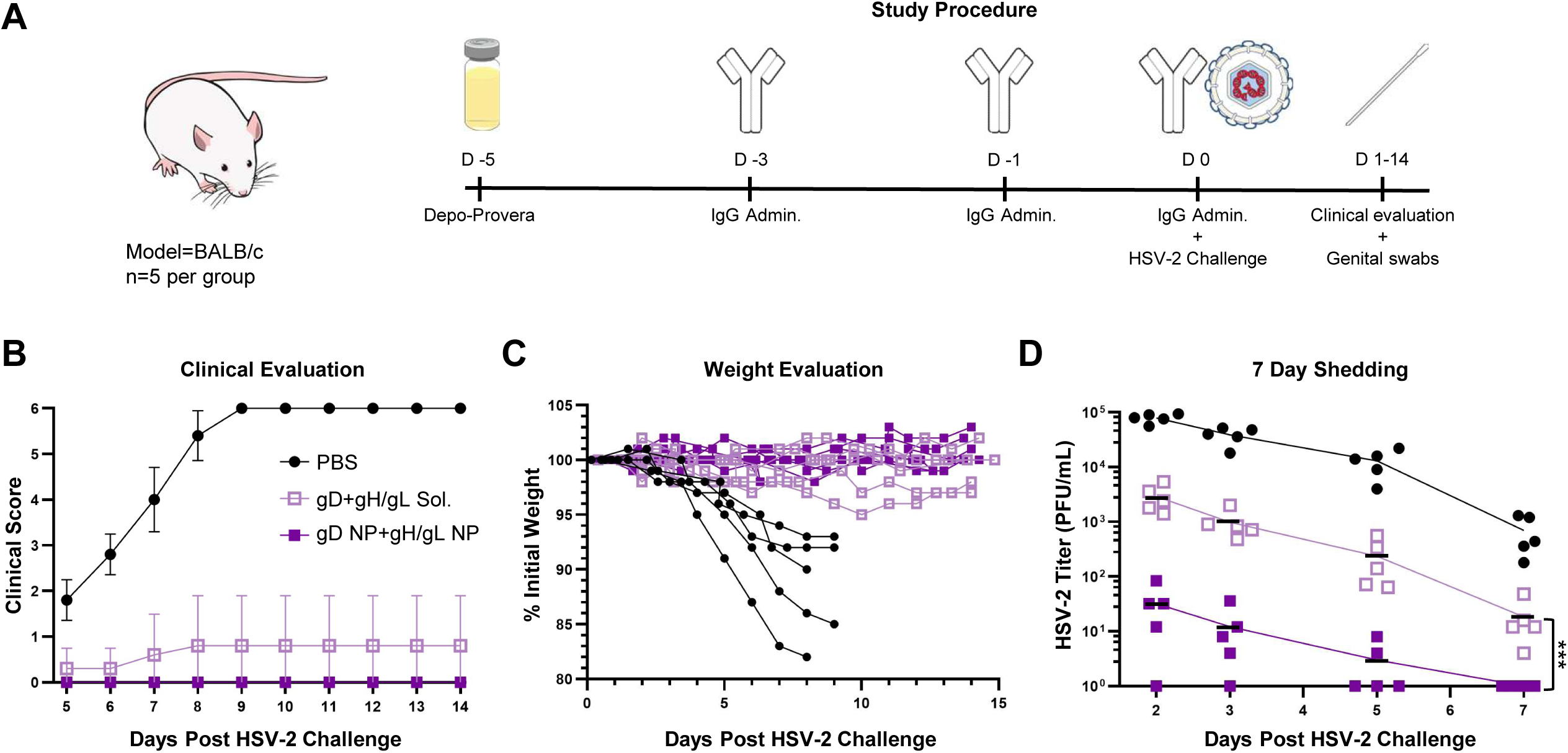
Protection after passive transfer of IgG from mice sera immunized with soluble gD+gH/gL, or gD NP+gH/gL NP from HSV-2 vaginal challenge. BALB/c mice (n=5 per group) were administered IgG intraperitoneally (IP) at 40 µg/gram of body weight on days -3, -1, and 0 of challenge. Mice were challenged intravaginally with 64,000 PFU of HSV-2 strain 333. (A) Study procedure of the passive transfer experiment. (B) Mice were monitored daily for clinical signs of HSV-2 infection including vaginal erythema, genital lesions, genital hair loss, ruffled fur, lethargy, abnormal gait, hunched back, or hind-limb paralysis and given a score of 0 (no disease) to 6 (dead) (see Methods). Data points indicate the means and error bars indicate standard deviations. The difference between gD+gH/gL Sol. and gD NP+gH/gL NP was not statistically significant. (C) Weight was taken daily for 2 weeks post challenge and data points indicate individual mice. Mice receiving PBS did not survive past day 9. The differences between gD+gH/gL Sol. and gD NP+gH/gL NP were not statistically significant. (D) HSV-2 titers from vaginal swabs taken on days 2, 3, 5, and 7 post-challenge with HSV-2. Data points indicate individual mice and means are shown as bars. ***P□<□0.001.

Passive immunization resulted in no detectable clinical disease in mice receiving IgG from gD NP+gH/gL NP vaccinated mice, while 2 of 5 mice receiving IgG from soluble gD+gH/gL vaccinated mice developed vaginal erythema and lesions (**Figure 5B**). All PBS control mice died by day 9. Mice were also weighed daily for two weeks after challenge (**Figure 5C**). By day 7, all PBS control mice were under 95% of their original weight, and by their deaths, mice had 82%-93% of their original weight. All 5 mice receiving IgG from gD NP+gH/gL NP vaccinated mice lost no weight, while 3 mice receiving IgG from soluble gD+gH/gL vaccinated mice had no weight loss and 2 mice reached to 95% and 97% of their pre-challenge weight on day 10, but then recovered to 97% and 98%, respectively, by day 14. Mouse vaginal tracts were swabbed on days 2, 3, 5, and 7 post-challenge to quantify levels of infectious virus (**Figure 5D**). On day 2, PBS control mice had the highest mean titers at 79,200 PFU/mL, followed by mice receiving IgG from soluble gD+gH/gL vaccinated mice (mean=2,920 PFU/mL) and then mice receiving IgG from gD NP+gH/gL NP vaccinated mice (mean=32 PFU/mL). On day 5, only 2 of 5 mice receiving IgG from gD NP+gH/gL NP vaccinated mice had detectable virus (mean=2.4 PFU/mL), while all 5 mice receiving IgG from soluble gD+gH/gL vaccinated mice had infectious virus (mean=235.2 PFU/mL). Over the 7 day time course, mice passively immunized with IgG from gD NP+gH/gL NP vaccinated mice had lower virus shedding than mice receiving soluble gD+gH/gL (p<0.0001).

Overall, these data suggest that the IgG elicited by gD NP+gH/gL NP immunization is the major component responsible for the protection from HSV-2, and those antibodies are more protective than the response induced by immunization with soluble gD+gH/gL.

### gD NP+gH/gL NP Preferentially Elicit Antibodies Targeting Neutralizing Epitopes

Despite comparable binding antibody titers between the NP and soluble immunization groups (**Figure 2B, 2C, 4D, 4E**), NP immunization elicited markedly higher neutralizing antibody responses (**Figure 2D, 4C**) indicating improved functional antibody quality rather than increased overall antibody quantity. We hypothesized that immunization with HSV-2 NPs focused antibody responses to antigen domains critical for virus attachment compared with immunization with soluble antigens (**Supplemental Figure 6**). To test this hypothesis, we used a panel of mouse anti-gD monoclonal antibodies (mAbs) that ranged from non-neutralizing to potently neutralizing, including mAbs that target specific gD receptor binding domains. We performed serum competition assays using biolayer interferometry to determine which mAbs could compete with antibodies in the serum of mice immunized with gD NP+gH/gL NPs or soluble gD+gH/gL for binding to gD. We used 4 anti-gD mAbs in this experiment: MC14 (non-neutralizing), MC16 (non-neutralizing), DL11 (neutralizing, targets nectin-1 and HVEM binding sites), and MC23 (neutralizing, targets nectin-1 binding site)^27,44^. Serum from mice immunized with gD NP+gH/gL NP had higher levels of antibody that blocked neutralizing mAbs targeting the receptor binding domains of gD (nectin-1 and HVEM) compared to serum from mice immunized with soluble gD+gH/gL (DL11, 34.5% vs. 20.3%, respectively, p=0.042 and MC23, 41.2% vs. 27.2%, respectively, p=0.0002) **(Figure 6A, 6B)**. Furthermore, the range of blocking activity was highly variable in mice that received soluble gD+gH/gL, while responses by the gD NP+gH/gL NP group showed less variation. When non-neutralizing mAb MC14 was used as a competitor, serum from mice immunized with soluble gD+gH/gL had higher blocking activity than serum from mice immunized with gD NP+gH/gL NP (27.6% vs. 20.2%, respectively, p=0.037) **(Figure 6C)**. With non-neutralizing mAb MC16 as the competitor, blocking between soluble gD+gH/gL and gD NP+gH/gL NP was comparable (31.2% vs. 26.8%, p=0.49) **(Figure 6D)**. Taken together, these data suggest that the marked increase in neutralizing antibody titers elicited by gD NP+gH/gL NP compared with soluble gD+gH/gL is likely due to more focused targeting of antibodies to HSV-2 gD domains critical for binding to virus host cell receptors.

**Figure 6.**
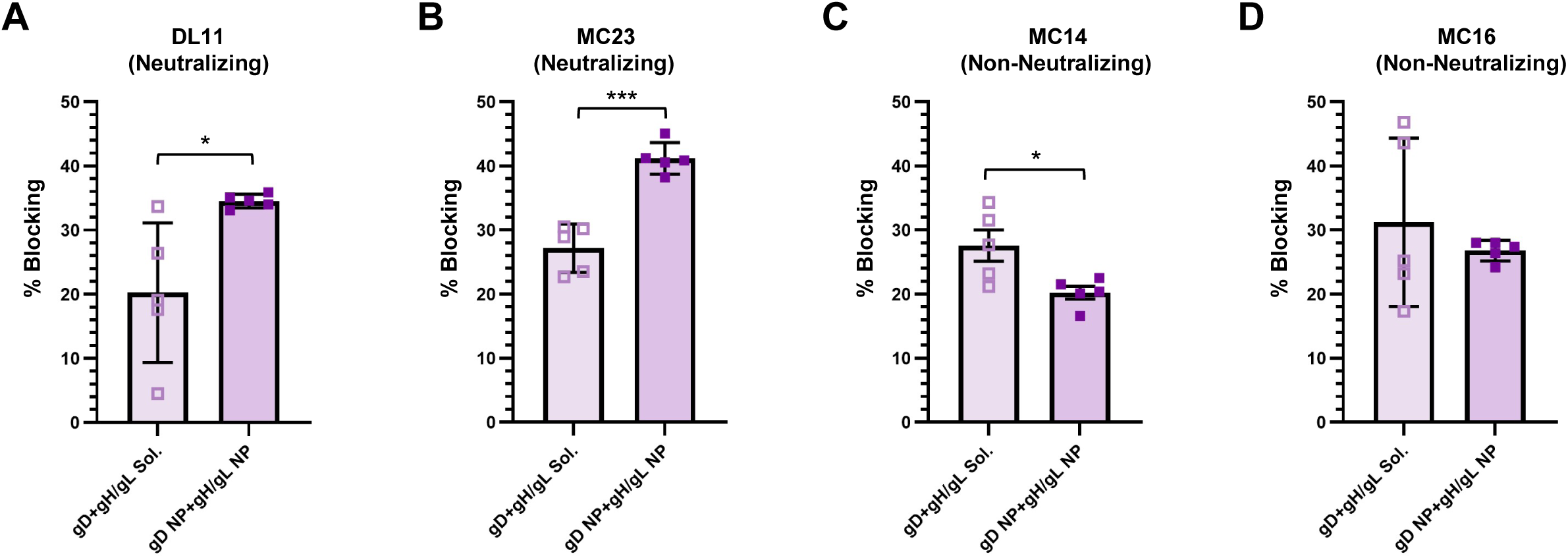
Detection of antibodies in sera from immunized mice that compete with known anti-HSV-2 gD neutralizing monoclonal antibodies. (A-D) Cross-competition of immune sera (n=5 per group) to neutralizing (A, B) or non-neutralizing (C, D) gD murine monoclonal antibodies. Bars indicate the means and error bars indicate standard deviations. For neutralizing antibodies, DL11 targets HVEM and nectin-1, and MC23 targets nectin-1. *P□<□0.05, ***P□<□0.001.

### HSV-2 Nanoparticles Induce High Titers of Neutralizing Antibodies and Antibodies that Block HSV-2 Glycoprotein Mediated Fusion in Non-Human Primates

Several HSV-2 vaccine candidates have demonstrated strong immunogenicity and efficacy in preclinical rodent models but have failed to confer protection in clinical trials^15^. Mice are not known to be naturally infected by alphaherpesviruses like HSV, but rhesus macaques are infected with an alphaherpesvirus- Cercopithecine herpesvirus 1 (B virus). Thus, the immune system of macaques has evolved with a virus similar to HSV^45^. To evaluate vaccine-induced antibody responses in a more closely related species to humans, we immunized macaques with 50 µg of soluble gD+gH/gL or gD NP+gH/gL NP at an equimolar ratio of gD to gH/gL formulated with SAS at weeks 0 and 4 (**Figure 7A**). Two doses of vaccine were given IM 4 weeks apart and animals were bled 4 weeks post-dose 1 and 2-weeks post-dose 2 (**Figure 7A**). Macaques immunized with gD NP+gH/gL NP had higher anti-gD binding antibody titers than those vaccinated with soluble gD+gH/gL after each dose (dose 1, 10,400 vs. 300, respectively, p=0.046; dose 2, 115,200 vs. 15,400, respectively, p=0.037) (**Figure 7B**). On the contrary, anti-gH/gL antibody titers between gD NP+gH/gL NP and soluble gD+gH/gL were comparable after each dose (dose 1, 2,000 vs. 1,400, respectively; dose 2, 86,400 vs. 48,000, respectively) (**Figure 7C**).

**Figure 7.**
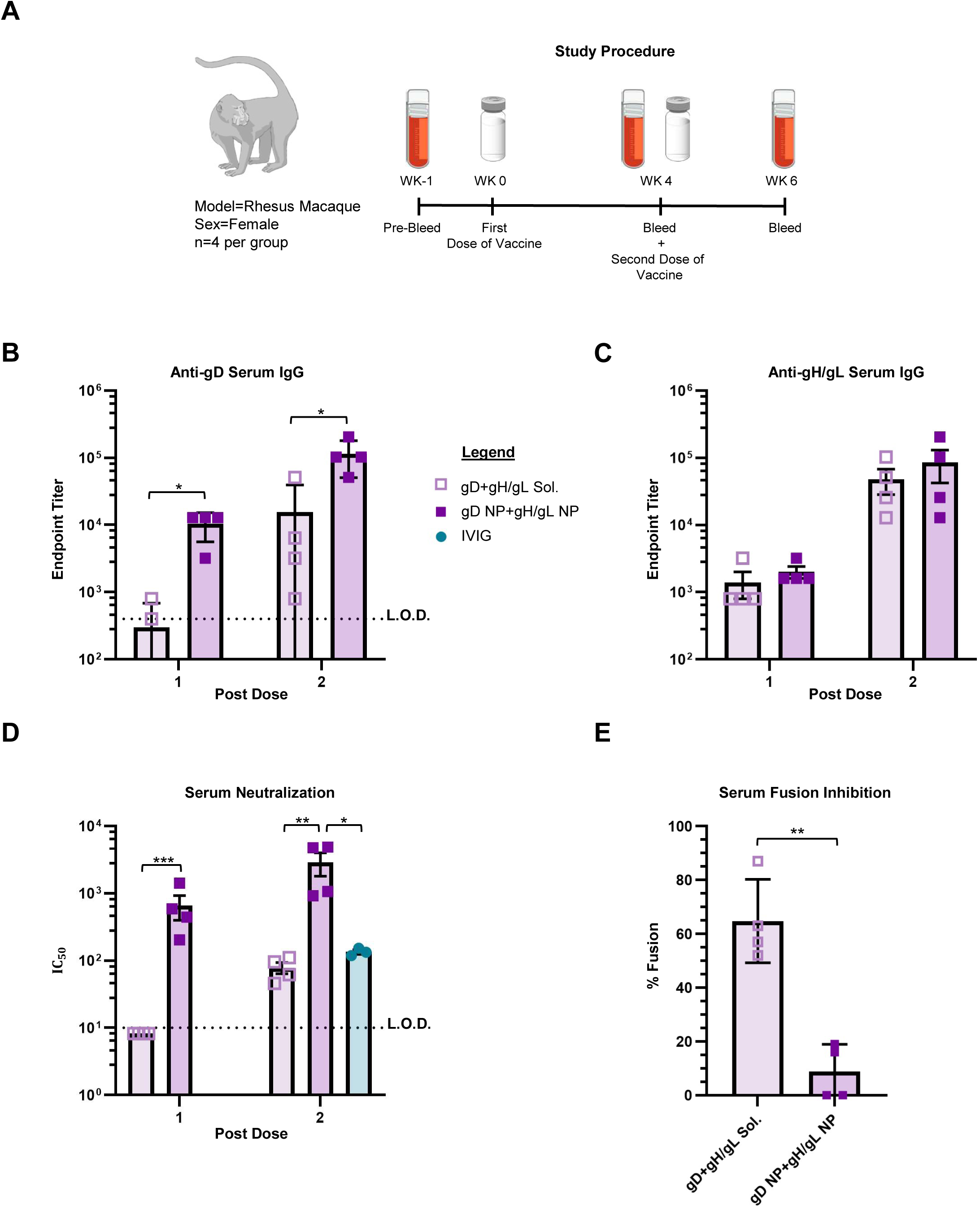
Antibody responses in rhesus macaques immunized with gD+gH/gL Sol. or gD NP+gH/gL NP. (A) Schematic of the study. Rhesus macaques (n=4 per group) were immunized intramuscularly with 2 doses of vaccines at 50 µg per dose in SAS adjuvant given 4 weeks apart. Serum was analyzed from macaques bled 4 weeks after the first dose and 2 weeks after the second dose of vaccine. (B) HSV-2 gD binding ELISA antibody titers in serum shown as endpoint titers. L.O.D. is limit of detection defined as an endpoint titer of 1:400. Bars indicate means and error bars indicate standard deviations. *p<0.05. (C) HSV-2 gH/gL binding ELISA antibody titers in serum shown as endpoint titers. Bars indicate means and error bars indicate standard deviations. Differences were not statistically significant. (D) HSV-2 neutralizing antibody titers in serum and comparison to human IVIG (diluted 9-fold so that the concentration of total IgG is the same as in humans). IC_50_ is the dilution of sera that inhibits infection by 50%. Bars indicate means and error bars indicate standard deviations. L.O.D. is the limit of detection defined as an IC_50_ of 10. * p <0.0167; ** p<0.0033; *** p<0.0003. (E) HSV-2 fusion inhibition from antibody responses in serum. Bars indicate means and error bars indicate standard deviations. Percent fusion was defined as the luminescence of immune serum samples divided by the luminescence of pooled pre-immune serum samples x 100. ** p<0.01.

After only 1 dose, macaques immunized with gD NP+gH/gL NP had mean neutralizing antibody titers at 664.5, while neutralizing antibodies were not detected in animals vaccinated with soluble gD+gH/gL (p=0.0006) (**Figure 7D**). After dose 2, neutralizing antibody titers were boosted in macaques vaccinated with gD NP+gH/gL NP, and the level of neutralizing antibody titers were ∼37-fold higher than macaques immunized with soluble gD+gH/gL (2,900 vs. 78.8, respectively, p=0.0022) **(Figure 7D)**. To provide a contextual benchmark for the neutralizing antibody titers elicited in immunized macaques, we compared these responses to human intravenous immunoglobulin (IVIG). IVIG is derived from plasma pooled from >1,000 human donors and represents the aggregate antibody repertoire present in the population from the country in which it is manufactured, including individuals naturally infected with HSV. In the United States, among persons aged 14-49, the estimates of seroprevalence for HSV-1 is 47.8% and for HSV-2, 11.9%^46^. Therefore, neutralization assays with IVIG offer a practical surrogate for average human HSV-specific humoral immunity, enabling a rough comparison of neutralizing antibody magnitude. Compared to IVIG (when diluted 9-fold so that the total IgG is like that in humans), macaques immunized with gD NP+gH/gL NP had higher neutralizing antibody titers (p=0.0079) and macaques immunized with soluble gD+gH/gL had comparable titers (p=0.055).

Since our vaccine strategy targets both virus attachment (gD) and virus fusion to the host cell (gH/gL), we evaluated whether serum from immunized macaques could also inhibit HSV-2 fusion. Serum from macaques immunized twice with gD NP+gH/gL NP inhibited fusion ∼7-fold more effectively compared to serum from macaques immunized with soluble gD+gH/gL (8.8% vs. 64.76%, p=0.0016) (**Figure 7E**).

In summary, these data indicate that in non-human primates, immunization with gD NP+gH/gL NP elicit higher HSV-2 neutralizing and fusion blocking antibody responses than immunization with soluble gD+gH/gL. In addition, the HSV-2 neutralizing antibody responses in non-human primates elicited by gD NP+gH/gL NP were higher than those seen in natural HSV-2 infection of humans.

## Discussion

HSV-2 directly infects mucosal surfaces, uses a complex viral entry process and has multiple immune evasion strategies, which has made creating a successful vaccine daunting^12^. An effective HSV-2 vaccine is likely to require optimizing gD as an immunogen, along with targeting of other glycoproteins that are essential for virus entry^29^. Here, we show that vaccination with NPs separately displaying gD and gH/gL induce potent immune responses in mice and non-human primates and are highly effective in protecting mice from HSV-2 challenge.

The largest phase 3 clinical trial of an HSV-2 prophylactic vaccine used soluble gD adjuvanted with alum and 3-O-deacylated monophosphoryl lipid A, but failed to protect against HSV-2 infection or disease^31^. One possible explanation for the lack of success in the trial is the relatively low and short-lived neutralizing antibody titers induced by the vaccine, along with low or absent antibody responses to several critical gD epitopes^27,47^. Arrayed multivalent NP display of antigens can increase both the magnitude and the durability of neutralizing antibody responses, and neutralizing antibody titers can be enhanced through rational design with epitope-focusing strategies^34,39^. Hence multivalent display is a promising approach in enhancing gD as a protective immunogen.

Another possible explanation for the lack of success of the phase 3 trial was the use of a single glycoprotein as an immunogen. Additional viral glycoproteins are required for HSV-2 entry, including gH/gL. HSV-2 gH/gL forms a boot-like shape^16^ which is bulkier than the rod-like shaped gH/gL present in Epstein-Barr virus (EBV),^16^ and when HSV-2 gH/gL is genetically fused to the nanoparticle scaffold, steric hinderance may become an obstacle to NP display^48,49^. Attempts to design an HSV-2 gH/gL ferritin NP vaccine have been unsuccessful in our laboratory (data not shown). Since SpyTag/SpyCatcher coupling to NPs has been validated for high molecular weight and complex antigens of varying stability and symmetry (e.g. HIV envelope, influenza hemagglutinin and neuraminidase)^41,50^, we used this approach and successfully displayed HSV-2 gH/gL on a NP (**Figure 1**).

Immunization of mice with 2 doses of gD NP or gH/gL NP elicited higher titers of HSV-2 neutralizing antibodies in comparison to the corresponding soluble glycoproteins, although anti-gD and anti-gH/gL binding antibody levels were comparable. This suggested that the immune response was focused towards more neutralizing epitopes on the target glycoproteins. Prior studies with NPs showed that orientation of the antigen on the particle can result in preferential induction of antibodies to specific epitopes^39,51^. Serum competition assays using neutralizing mAbs indicated that serum from mice immunized with gD NP+gH/gL NP had more antibodies to HSV-2 gD neutralizing epitopes than mice immunized with soluble gD+gH/gL (**Figure 6**).

While neutralizing antibodies that block virus attachment to cells are likely important for protection from infection, our data indicate that antibody-mediated protection against HSV-2 is not fully explained by conventional neutralizing activity as measured by the standard Vero-cell based neutralization assay. Passive transfer experiments (**Figure 5**) demonstrate that vaccine-elicited IgG is sufficient to confer protection in vivo, establishing a central role for humoral immunity. However, this protection does not correlate strictly with serum neutralizing titers, as antibodies targeting gH/gL exhibit strong fusion-blocking activity yet, when compared to serum from gD immunized mice, elicited lower neutralization in standard in vitro assays (**Figure 2**, **Figure 3**). This apparent discordance suggests that commonly used neutralization assays, which are optimized to detect inhibition of early viral entry events such as receptor engagement, may underrepresent antibody functions that act at later stages, including membrane fusion. In addition, these assays do not capture other downstream antibody activities such as inhibition of cell-to-cell spread^52^ or post-attachment neutralization^53^, which can contribute to antiviral immunity. Antibodies directly targeting fusion have implicated in protection for EBV^48^. Together, these findings support a model in which antibodies elicited by multivalent NP vaccination mediate protection through complementary mechanisms, including but not limited to classical neutralization, and highlight the limitations of relying solely on in vitro neutralizing antibody titers as a correlate of protective immunity.

We postulated that vaccination with a combination of gD NPs and gH/gL NPs would provide better protection than vaccination with either alone, since gD and gH/gL mediate different steps of viral entry^16,22,23^. Antibodies that target gD can block receptor binding domains^27^, while anti-gD-gH/gL antibodies block virus fusion^54^. Immunization with 2 doses of gD NP+gH/gL NP followed by challenge 2 weeks later with a lethal dose of HSV-2 nearly eliminated shedding and prevented virus entry into dorsal root ganglia, which suggests that the vaccine might prevent establishment of HSV-2 latency (**Figure 3**). Of interest, we found that immunization with gD NP+gH/gL NP was more effective than with gD NPs or gH/gL NPs alone but combining soluble gD and gH/gL had little to no added protection compared to soluble gD alone (**Figure 3**). These results are consistent with a prior study which found that that adding soluble gH/gL and soluble gB to a soluble gD vaccine provided no benefit^33^.

An important goal for an effective HSV-2 vaccine is to provide durable protective immunity. In the large phase 3 HSV-2 vaccine trial evaluating soluble gD in HSV-2 seronegative women, the durability of the immune response was a major limiting factor; 5 months after volunteers received 2 doses of vaccine, median neutralizing antibodies were undectable^31^. The durability of the humoral immune responses correlates with prolonged germinal center B cell activity and enhanced somatic hypermutation, mechanisms implicated in the success of human papillomavirus VLP vaccines^55,56^. We found that neutralizing antibody titers elicited by gD NP+gH/gL NP persist longer and at higher titers than responses generated by soluble gD+gH/gL (**Figure 4**).

Compared with mice, non-human primates are genetically more similar to humans and some HSV immune evasion proteins are functional in non-human primate cells^57^, but not mouse cells^58^. HSV-2 is thought to have evolved by transmission from an ancestor of non-human primates to a precursor of modern humans^59^. Thus, HSV-2 has co-evolved with non-human primates and immune responses to HSV-2 vaccines in these animals may be more predictive of responses in humans than immune responses in mice. Therefore, it is encouraging that all of the macaques had robust antibody responses; 2 doses of gD NP+gH/gL NP induced 37-fold higher titers of HSV-2 neutralizing antibody in macaques than soluble gD+gH/gL (**Figure 7D**). Sera from macaques immunized with gD NP+gH/gL NP had 7-fold higher fusion-blocking activity compared to soluble gD+gH/gL (**Figure 7E**). Furthermore, HSV-2 neutralizing antibody titers in macaques were 22-fold higher than titers in plasma from HSV-2 infected humans (**Figure 7D**). While it is difficult to compare the absolute values of HSV-2 neutralizing antibody titers among different laboratories, a study using a combination vaccine containing HSV-2 soluble gD, gE, and gC in macaques showed that after 3 doses of the vaccine, HSV-2 neutralizing antibody titers in monkeys (∼160) were similar to titers seen in natural infected humans (∼160)^60^. Thus, the higher HSV-2 neutralizing antibody titers with the gD NP+gH/gL NP vaccine in macaques compared to those in naturally infected humans suggests that gD NP+gH/gL NP is a promising vaccine candidate.

Our study has some limitations. Although gD NP+gH/gL NP fully protected from disease and markedly reduced shedding, HSV-2 vaccine studies in mice may not be predictive of vaccine efficacy in humans^15^. While competition assays gave more insight into the mechanism of protection conferred by gD NP+gH/gL NP, the mAbs evaluated were only against one component of the vaccine, gD. More anti-HSV-2 gH/gL mAbs must be isolated and characterized to better understand the mechanism of the superior protection granted by gH/gL NP immunization. Furthermore, although we were able to evaluate immunogenicity in macaques, we could not evaluate protection from HSV-2 challenge, since macaques shed little or no virus and do not develop genital lesions after infection with HSV-2^60,61^.

In summary, we found that a vaccine combining nanoparticles that display HSV-2 gD and gH/gL was highly protective against disease, shedding, and infection of dorsal root ganglia in mice challenged with a lethal dose of HSV-2. Immunization with gD NP+gH/gL NP elicits high level and durable antibody titers, with immune responses focused towards neutralizing epitopes on HSV-2 gD. Passive transfer shows that IgG elicited by gD NP+gH/gL NP is the principal component for protection from HSV-2 in mice. The discordance between neutralization titers and protection conferred by gD NP vs. gH/gL NP suggested that the standard neutralization assay incompletely captures the importance of fusion-dependent inhibition. Mice and monkeys receiving gD NP+gH/gL NP develop high titers of neutralizing and fusion-blocking antibodies. Therefore, the rationally designed gD NP+gH/gL NP vaccine is a promising vaccine candidate to reduce HSV-2 infection and/or disease.

## Methods

### Plasmids and Cloning

Plasmids were cloned in *Escherichia coli* DH5α cells using standard PCR protocols with Q5 High-Fidelity 2× Master Mix (New England Biolabs) and Gibson Cloning. Plasmids were validated by Sanger Sequencing. pET28a-SpyCatcher003-mi3 (GenBank MT945417, Addgene 159995) and pDEST14-SpySwitch (GenBank ON131074, Addgene 184225) were previously described^40,41^. HSV-2 gD, gH, and gL were isolated from HSV-2 strain 333^62^. For gD333ST3, a cassette containing HSV-2 gD (amino acids 1-333) followed by an 11 amino acid linker (GSGGSGGSGTG) and a 16 amino acid SpyTag (RGVPHIVMVDAYKRYK) was inserted into pcDNA3.1. For gH803ST3gL, A cassette containing HSV-2 gH (1-803) followed by the same linker and SpyTag used for HSV-2 gD was inserted into pVRC8400. Full length HSV-2 gL was inserted into plasmid pVRC8400 (Addgene).

### Expression and Purification of SpyCatcher003-mi3 Nanoparticles

SpyCatcher003-mi3 was expressed in *E. coli* BL21(DE3) transformed with pET28a-SpyCatcher003-mi3 and plated on an LB-agar plate containing 50 μg/mL kanamycin. A single colony was used to inoculate LB medium supplemented with kanamycin, grown overnight, and then expanded into 1 L of LB with kanamycin and incubated at 37°C until the OD600 reached 0.6, at which point IPTG was added to a final concentration of 0.5 mM. The culture was then incubated at 22°C for 16 h and then pelleted by centrifugation at 4,000 x g.

Cell pellets were resuspended in 20 mL of 20 mM Tris-HCl (pH 8.5), 300 mM NaCl containing 0.1 mg/mL lysozyme, 1 mg/mL protease inhibitor (cOmplete, EDTA-free; Roche), and 1 mM phenylmethylsulfonyl fluoride. After incubating at 4°C for 45 min with mixing, cells were lysed by sonication on ice (4 cycles, 60 seconds per cycle, 40% duty cycle, Q2000 Sonicator (QSONICA). The lysate was clarified by centrifugation at 35,000 x g for 45 min at 4°C. To precipitate nanoparticles, 170 mg of ammonium sulfate was added per mL of supernatant, followed by incubation at 4°C for 1 hr with mixing. The precipitate was collected by centrifugation at 30,000 x g for 30 min at 4°C, and the pellet was resuspended in mi3 buffer (25 mM Tris-HCl, 150 mM NaCl, pH 8.0). The solution was filtered through 0.22 µm syringe filter before dialysis overnight (1,000-fold excess mi3 buffer). Dialyzed protein was further clarified by centrifugation at 17,000 x g for 30 min at 4°C and filtered again through a 0.22 µm filter. For purification, the sample was loaded onto a HiPrep Sephacryl S-400 HR 16/60 column (GE Healthcare) pre-equilibrated with mi3 buffer and separated at 0.1 mL/min using an ÄKTA Pure 25 system (GE Healthcare), collecting 1.5 mL fractions. Purified fractions were pooled and concentrated using a Vivaspin 20 (100 kDa MWCO) centrifugal concentrator before storage at −80°C.

### Expression and Purification of HSV-2 Glycoprotein Constructs

The gD333ST3 and gH803ST3gL constructs were expressed in Expi293F cells (Thermo Fisher Scientific). Cells were cultured in Expi293 Expression Medium (Thermo Fisher Scientific) at 37°C with 8% (v/v) CO□. Transfections were carried out using an ExpiFectamine 293 Transfection Kit (Thermo Fisher Scientific). Expi293F cells were adjusted to a density of 3 × 10 cells/mL, and 1 µg of plasmid DNA per mL of culture was incubated with ExpiFectamine 293 reagent for 15 min before being added dropwise to the cell culture. gH803ST003 and gL were co-transfected in a 1:1 molar ratio. After 16–20 hr, ExpiFectamine 293 Transfection Enhancers 1 and 2 were added. Cell supernatants were harvested 7 days post-transfection by centrifugation at 4,000 x g for 5 min at 4°C, and the remaining supernatant was filtered through a 0.22 µm filter to remove cell debris.

Purification of gD333ST003 and gH803ST003gL was performed using the SpySwitch system at 4°C^40^. SpySwitch buffer (50 mM Tris-HCl, pH 7.5, 300 mM NaCl) at 10x concentration was added to the mammalian culture supernatant at a final concentration of 10% (v/v). SpySwitch resin, packed into an Econo-Pac Chromatography Column (Bio-Rad), was pre-equilibrated with 20 column volumes (CV) of SpySwitch buffer^40^. The culture supernatant was then loaded onto the column at a low flow rate using a peristaltic pump, and the flow-through was collected. Following sample loading, the column was washed twice with 15 CV of SpySwitch buffer. Elution was performed by incubating the column with 1.5 CV of SpySwitch Elution Buffer (50 mM acetic acid/sodium acetate, pH 5.0, 150 mM NaCl) at 4°C with the column capped. After incubation for 5 min, the cap was removed, and the elution flow-through was collected directly into a microcentrifuge tube containing 0.3 CV of 1 M Tris-HCl (pH 8.0) and the tube was gently mixed by inversion to minimize time of exposure to low pH. This elution step was performed six times to maximize protein recovery. Typical yields for gD333ST003 were 70-140 mg/L of culture, and typical yields for gH803ST003gL were 30-50 mg/L of culture. Purity was assessed by SDS–PAGE followed by Coomassie staining. Gel images in the figures were given a blue color from the original black and white images. Purified proteins were aliquoted and stored at −80°C.

### Endotoxin depletion and quantification

Endotoxin was removed from all vaccine components using the Triton X-114 phase separation method^63^. Protein samples were incubated on ice with Triton X-114 at a final concentration of 1% (v/v) for 5 min. The mixture was then incubated at 37°C for 5 min, followed by centrifugation at 16,000 x g for 1 min at 37°C. The aqueous phase was transferred to a fresh tube. This process was repeated three times, and a final wash without Triton X-114 was performed to remove any residual detergent. To quantify endotoxin levels, a Pierce Chromogenic Endotoxin Quant Kit (Thermo Fisher Scientific) was used following the manufacturer’s instructions.

### Conjugation of Immunogens onto Nanoparticles

The concentration of vaccine components was measured using a Bradford Assay (Pierce). To make the conjugated nanoparticles, Spy-tagged antigens were incubated with unconjugated SpyCatcher003-mi3 at a 1.5 (gD) or 3 times (gH/gL) molar excess of antigens to mi3 monomers for 16 hr in TBS (50 mM Tris-HCl, 150 mM NaCl, pH 8.0) at 4°C. Excess antigen was removed with size-exclusion chromatography. Nanoparticles were loaded onto a HiPrep Sephacryl S-500 HR 16-600 column (GE Healthcare), which was equilibrated with PBS pH 7.4 and run on an ÄKTA Pure 25 system (GE Healthcare). The proteins were separated at 0.5 mL/min while collecting 1 mL elution fractions. Particles were concentrated and then assessed by SDS-PAGE with Coomassie blue staining to confirm separation of excess unconjugated antigen and conjugated nanoparticle. Particle homogeneity was assessed using dynamic light scattering. Briefly, NPs were centrifuged at 16,900g at 4°C for 30 min and then incubated at 20°C before readings. 5 µL of the supernatant was loaded into a quartz cuvette and samples were measured at 20°C using a NanoStar II (Wyatt Technology) with 10 scans of 5 s each. The intensity of the size distribution was averaged over 10 runs and normalized to the peak value using Microsoft Excel and the mean hydrodynamic radius and polydispersion values were shown.

### Negative Stain Electron Microscopy

3 μL of sample was applied to glow-discharged 200-mesh copper grids with a carbon support film (Ted Pella, Redding, CA). After settling for 30 s, the sample was wicked from the grid and briefly washed with distilled H_2_O. Grids were then negatively stained for 30 s with Nano-Van stain, consisting of 2% methylamine vanadate (Nanoprobes, Yaphank, NY). Excess stain was wicked from grids, and grids were allowed to air dry before imaging. Grids were imaged at 120 kV with a Talos transmission electron microscope using a bottom mount-Ceta digital camera (Thermo Fisher Scientific).

### Cryogenic Electron Microscopy (CryoEM) Grid Preparation and Data Collection

3 µL of samples were applied to freshly glow-discharged C-flat grids (Protochips) using an easiGlow system. Grids were blotted for 2.5 sec at 6°C and 100% humidity with a blot force of 4 pN using a Vitrobot Mark IV (Thermo Fisher Scientific), then plunge-frozen in liquid ethane and stored in liquid nitrogen. CryoEM data were collected on a Glacios transmission electron microscope (Thermo Fischer Scientific) operating at 200kV and a nominal magnification of 150,000x, using a Falcon 4i direct electron detector in counting mode. SerialEM was used for data collection.

### CryoEM Data Processing

Patch motion correction and Patch Contrast Transfer Function (CTF) estimation were carried out in cryoSPARC 4.7.1 on 6,472 movies. Particles from blob picking were used to create 2D templates after several rounds of 2D classification. These high-quality templates were then used for re-picking particles. Multiple rounds of 2D classification helped improve data quality. *Ab initio* reconstruction with 3 classes, followed by heterogeneous refinement, was used to select the best class. Two rounds of global and local CTF refinement, reference motion correction, and non-uniform refinement were conducted to produce the final reconstruction at 3.42 Å resolution from 78,161 particles (**Supplemental Figure 4**). Icosahedral symmetry was applied during refinement. A monomer structure from PDB 7B3Y was docked into the cryoEM map using ChimeraX and the whole nanoparticle structure was generated through icosahedral symmetry.

### Mouse Immunization, blood sampling, and HSV-2 challenge

All mouse studies were performed under a protocol approved by the Animal Care and Use Committee of the National Institute of Allergy and Infectious Diseases. Female BALB/c mice were purchased from Envigo RMS Inc. and were inoculated at 6 weeks of age. 5 mice per group were immunized intramuscularly with 5 µg of total soluble protein or nanoparticle normalized to the mass of antigen with SAS adjuvant (Sigma-Aldrich, 1:1 [vol/vol]) for a total volume of 100 µL. Immunizations were performed 3 weeks apart for 2 doses. Mice were bled 1.5 to 2 weeks post immunization and sera were collected for further analysis. In the durability study, mice were bled 1, 5, and 8 months post dose 2.

Mice were challenged 2 weeks after the second immunization and 9 months after the second immunization in the durability study. To thin the vaginal lining and synchronize the menstrual cycle, mice were injected subcutaneously with 2 mg of medroxyprogesterone acetate diluted in phosphate-buffered saline (PBS) 5 days before immunization. Mice were challenged intravaginally with 10 µL of 64,000 PFU (63xLD_50_) of HSV-2 strain 333^62^. To detect virus replication in the vaginal tract, vaginal swabs were obtained on days 1-7 after challenge, placed in vials filled with 300 µL of Dulbecco’s Modified Eagle’s Medium (DMEM), and were stored at -80°C until further processing. Serial dilutions of the vaginal swab solution were added to Vero cell monolayers in 6-well plates (after the media had been removed), incubated for 1 hr at 37°C, the inoculum was removed, 2.5 mL of medium containing 0.3% (v/v) Gamunex-C (GRIFOLS) was added, and 2 days later plaques were counted as described above. Mice were monitored for survival and clinical signs for at least two weeks post challenge. On day 14 post-challenge, mice were euthanized and vertebral spinal segments were removed to harvest dorsal root ganglia (DRG)^64^.

Mice disease scores were determined on days 5 to 14 after HSV-2 challenge using the following scoring system: 1- perineum with mild erythema or some hair loss, 2- perineum with moderate erythema or hair loss or edema, 3- perineum with more than one of the following: erythema, edema, hair loss, 4- perineum with more than two of the following: erythema, edema, wet, hair loss; or mild systemic symptoms like hunched posture, mild hindlimb paresis, 5- perineum disease plus hindlimb paralysis or lethargy; 6, death. If mice reached a score of 5, they were humanely euthanized the following day.

### Murine Dorsal Root Ganglia (DRG) DNA Extraction

DRG from vertebral spinal segments L3 to S4 were collected in RPMI 1640. DNA was isolated using a QIAgen DNeasy Blood and Tissue Kit (QIAgen 69504) with some modifications. DRG tissues were minced with a scalpel and resuspended with 40 µL of proteinase K and 360 µL of ATL lysis buffer. The tubes were then incubated at 56°C for 2 hr and then vortexed for 5 min. The remainder of the procedure followed the protocol from the QIAgen kit; DNA was eluted with 200 µL of AE buffer. The DNA concentration was determined from absorbance at 260 nm.

### Quantification of HSV-2 DNA in DRG

The amount of HSV-2 DNA in murine DRG was quantified by Taqman real time quantitative polymerase chain reaction (qPCR) with primers and probes specific for HSV-2 gG gene^65^. Each PCR reaction volume totaled 25 µL, including 12.5 µL of 2x TaqMan Universal PCR Master Mix (ThermoFisher Scientific, 4304437), 10 µL of sample DNA, 1 µM of primers gG2F2 (5’ GCT CCC GCT AAG GAC ATG C 3’) and gG2R2 (5’ GAT GAT AAA GAG GAT ATC TAG AGC AGG G 3’), and 200 nM of probe gG2P2 (5’FAM-TCC CCC TGT TCT GGT TCC TAA CGG C-TAMRA3’). qPCR reactions were performed on a QuantStudio 3 instrument (Thermo Fisher Scientific, A28137) with the following reaction conditions: one cycle of incubation at 50°C for 2 min followed by 95°C incubation for 10 min and then 40 cycles of denaturation at 95°C for 20 sec and annealing at 60°C for 1 min. Analysis was done using QuantStudio 3 Design and Analysis Software v1.6.1 (Thermo Fisher Scientific). A standard curve was produced by plotting the qPCR amplification of 1:10 dilutions of HSV-2 gG plasmid DNA. All reactions were performed in duplicate, and the average of the duplicate values were plotted as HSV-2 DNA copy number per µg of total DNA isolated from each DRG tissue sample. None of the negative controls (PBS or ATL buffer/proteinase K alone) had detectable HSV-2 DNA by qPCR.

### Passive Transfer

6-week old BALB/c mice (Envigo, n=25) were immunized 2 times, 3 weeks apart, with PBS control, 5 µg of soluble gD+gH/gL, or 5 µg of gD NP+gH/gL NP with SAS adjuvant (Sigma-Aldrich, 1:1 (v/v)) for a total volume of 100 µL. 2 weeks after dose 2, mice were terminally bled and serum was collected and pooled and IgG was purified using a 5 mL protein G column (Cytiva). For passive transfer, 12-week old BALB/c mice were purchased from Envigo. Mice received medroxyprogesterone acetate and were challenged with the same protocol as explained above. On days -3, -1, and 0, mice received 40 ug of purified murine IgG diluted in sterile PBS per gram of body weight in a total of 100 uL administered intraperitoneally. Mice received purified IgG from mice previously given PBS or immunized with soluble gD+gH/gL or gD NP+gH/gL NP. Vaginal swabs were taken on days 2, 3, 5, and 7 post challenge and mice were followed for 2 weeks for evaluation of weight and disease.

### Non-Human Primate Immunization and Sampling

Non-human primate studies were performed under a protocol approved by the NIH Institutional Animal Care and Use Committee of the National Institute of Allergy and Infectious Diseases and were carried out in accordance with all federal regulations and NIH guidelines. Eight female herpes B virus-negative rhesus macaques, aged 7-12 years old, from a pathogen-free colony were used in this study and were seronegative for HSV-2 gD and gH/gL based on serum ELISA and serum neutralization titers. Macaques were immunized intramuscularly with 50 μg of soluble gD+gH/gL or gD NP+gH/gL NP (10 μg gD and 40 μg gH/gL) mixed 1:1 (vol/vol) with SAS adjuvant (Sigma-Aldrich) at weeks 0 and 4. Blood was drawn at weeks -1, 4, and 6 and sera was collected for analysis of gD and gH/gL binding assays and neutralizing antibody titers.

### HSV-2 Glycoprotein Binding Antibody

Immulon 4 HBX 96-well plates (Thermo Fisher Scientific) were coated with 1 µg/mL recombinant HSV-2 gD333ST3 or gH803ST3gL in PBS and incubated overnight at 4°C. Plates were then washed with PBSt (0.05% Tween in PBS) and blocked with a blocking buffer (2% bovine serum albumin in PBS) for 1 hr at room temperature. Sera were added to the plates in duplicate, diluted 3- to 5-fold and then incubated at room temperature for 1 hr. Plates were washed and incubated with HRP-conjugated secondary antibody specific for mouse IgG (Sigma-Aldrich, A9044) at a 1:3000 dilution for 1 hr. Plates were washed and developed with TMB substrate (KPL) for 5 min. The reaction was then stopped with 1 M H_2_S0_4_ and the 450 nm absorbance was measured using a spectrophotometer. Endpoint titers were calculated in Excel as absorbance greater than 2 times the mean absorbance of the background wells.

### HSV-2 Neutralization Assay

Vero cells were purchased from ATCC and maintained in DMEM containing 10% (vol/vol) fetal bovine serum (FBS), 100 U/mL penicillin, and 100 U/mL streptomycin at 37°C. Neutralizing antibody titers were quantified by adding serial dilutions of heat-inactivated (56°C for 30 min) mouse or rhesus macaque serum or IVIG (diluted 9x to obtain the same IgG level as found in human serum) in PBS to HSV-2 R519 (70 PFU, a plaque purified clone of HSV-2 333) and incubating for 1 hr at room temperature. After 1 hr, the serum-virus mixture (0.3 mL) was added to Vero cell monolayers in 6-well plates (from which the media had been removed), and incubated for 1 hr at 37°C. The inoculum was then removed, and DMEM (Gibco) supplemented with 10% (vol/vol) FBS and 0.3% (vol/vol) Gamunex-C (GRIFOLS) was added. The plates were then incubated at 37°C with 5% CO for 2 days, and then the cells were stained with a solution containing 10% (vol/vol) formaldehyde, 5% (vol/vol) acetic acid, 60% (vol/vol) methanol, and 1% (wt/vol) crystal violet for 30 min, washed, and plaques were counted. 50% plaque inhibition titers were calculated with GraphPad Prism (Software v.9.3.1), using nonlinear regression.

### HSV-2 Fusion Inhibition Assay

HSV-2 fusion was measured as previously reported^66^, with some modifications. Briefly, effector CHO-K1 cells were transfected with plasmids (0.48 µg each) encoding firefly luciferase under control of the T7 promoter, HSV-2 gB, gD, gH, and gL and 5 µL of Lipofectamine 2000 (Invitrogen) diluted in Opti-MEM (Invitrogen). CHO-K1 cells transfected with the same plasmids as the effector cells except without gB served as a negative control for fusion. Target CHO-K1 cells were transfected with plasmids expressing T7 RNA polymerase (0.48 µg) and nectin-1 (2 µg). Transfected cells were incubated overnight at 37°C. After transfection, effector or control cells were treated with trypsin, and 15,000 cells were plated onto wells of white bottom 96-well plates (Corning) and incubated for 30 min at 37°C with mouse or NHP pre-immune serum or serum post dose 2 at a 1:2 dilution in triplicate. After incubation, effector or control cells were mixed with target cells in a 1:1 ratio and incubated for 18 hr at 37°C. After the 18 hr incubation, luciferase activity was quantified using a luciferase reporter assay system (Promega) and a Varioskan LUX luminometer (Thermo Fisher Scientific). 100% fusion was defined as the average luminescence of the pooled pre-immune serum after subtraction of the background luminescence (wells with control cells, no gB added). For immune samples, background luminescence was subtracted from the average of the triplicates and % fusion was calculated as (immune luminescence/pooled pre-immune luminescence) x 100.

### HSV-2 Antibody Competition Studies

Antibody cross-competition was performed using biolayer interferometry. His-tagged gD protein expressed in Expi293 cells (1 μg/mL in PBS) was immobilized on Ni-NTA biosensors (Sartorius) for 300 s, followed by an equilibration step in PBS for 60 s. The biosensors were then dipped in competitor HSV-2 gD monoclonal antibodies (30 μg/mL in PBS) for 300 s, followed by another PBS equilibration step for 60 s. The biosensors were then dipped into wells containing serial dilutions of murine serum (five 2-fold dilutions starting at 1:40) for 300 s, followed by dipping into PBS for 180 s. Biolayer interferometry results were obtained using an Octet Red 96 instrument (FortéBio); assays were performed at 30°C with agitation at 1,000 rpm. The dilution factor was plotted against the corresponding maximum response unit and the area under the curve was calculated for each dilution series. Percentage competition was based on the area under the curve.

### Statistics

Statistics were performed on GraphPad Prism (Software v.9.4.1). Glycoprotein binding antibody titers, neutralizing antibody titers, and viral shedding data were log_10_ transformed for normalization purposes. For glycoprotein binding antibody titers, neutralizing antibody titers and fusion inhibition, the group averages were compared using the Welch Two Sample t test for each pair of groups followed with Bonferroni correction for multiple comparisons. The area under curve for virus shedding and clinical scoring was used to compare between groups. Statistical differences were analyzed using the Welch’s t-test when only two groups were compared, and when more than two groups were compared, the Welch Two Sample t-test was corrected for multiple comparisons using the Bonferroni correction. Specifically, p values less than 0.05/N (where N represents the number of multiple comparisons) were considered significant.

## Acknowledgements

D.M. received funding from the NIH OxCam program and Cambridge Trust. This research was supported in part by the Intramural Research Program of the National Institutes of Health (NIH). The contributions of the NIH author(s) are considered works of the United States Government. The findings and conclusions presented in this paper are those of the authors and do not necessarily reflect the views of the NIH or the U.S. Department of Health and Human Services. We thank Gary Cohen of the University of Pennsylvania for mouse monoclonal antibodies to HSV-2 gD, Richard Longnecker of Northwestern University for plasmids to the HSV-2 fusion assay, Jing Qin of NIAID for support in statistical evaluation of the data, and Camilla Schemmel of the University of Cambridge for support with figure design. Figure 1A and Supplemental Figure 6 were created by Ryan Kissinger from the Visual and Medical Arts Department with the Research Technologies Branch at NIAID. Illustrations found in figures 2, 4, 5, and 7 were created by the NIAID BioArt Source (https://bioart.niaid.nih.gov). This work utilized the computational resources of the NIH HPC Biowulf cluster (https://hpc.nih.gov). NIH Animal facility technicians supported mouse and non-human primate procedures.

## Contributions

D.M., A.V., G.L., and K.C.D. carried out mouse handling and documenting. K.K. carried out NHP sampling and immunizations and R.H. oversaw the NHP study. S.A.S. and F.H. carried out negative stain EM. H.L. and W.B. performed Cryo-EM data collection. D.M. and J.Z. carried out Cryo-EM data processing. D.M. and I.S.P. performed PCRs. D.M. carried out all other experiments. D.M., K.W., M.R.H., and J.I.C. designed the project. K.W., W.B., J.T.S., M.R.H, and J.I.C. provided critical guidance to the project. D.M., K.C.D, M.R.H., and J.I.C. wrote the manuscript. All authors read and approved the manuscript.

## Supplemental Figures

**Supplemental Figure 1.**
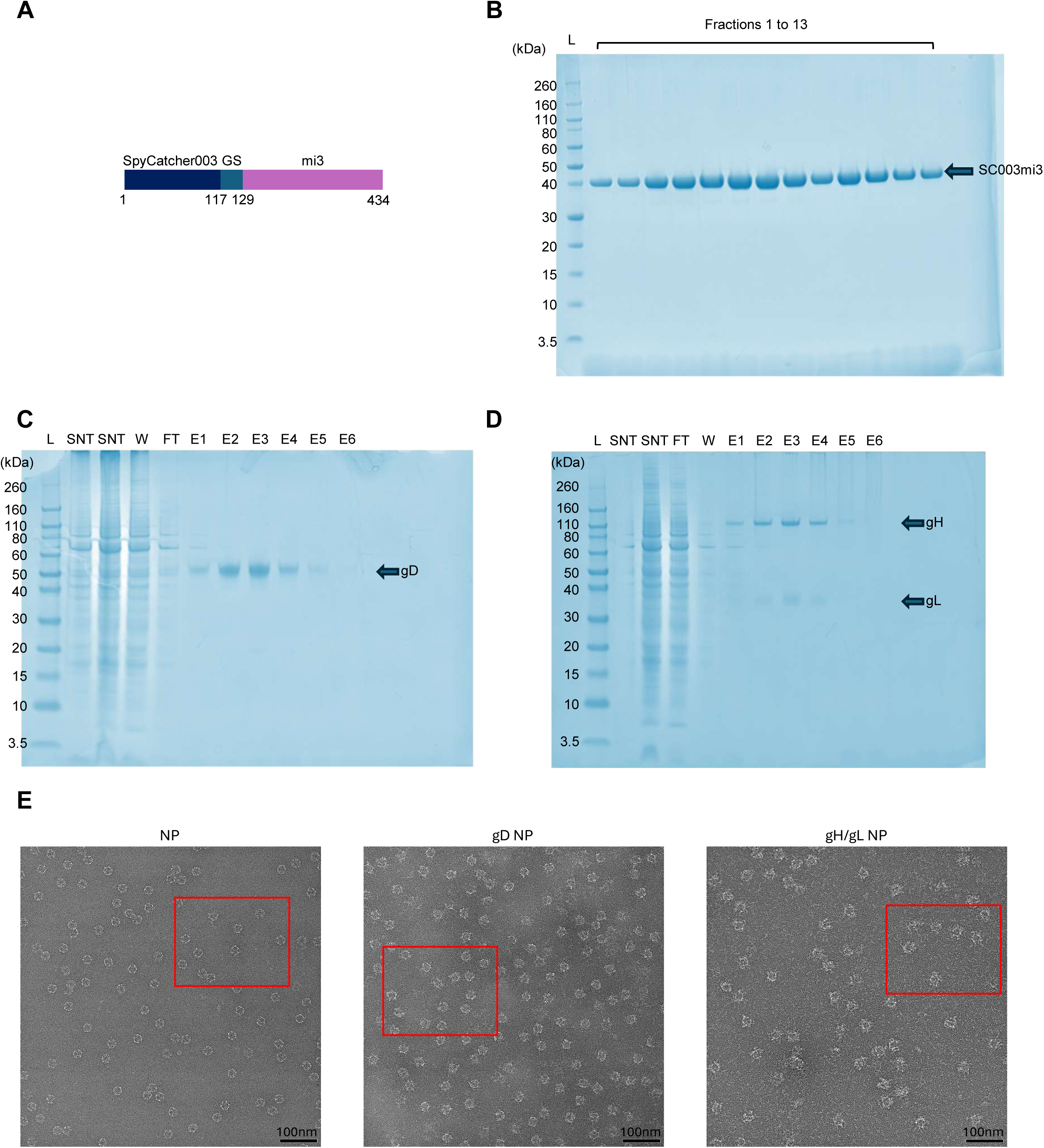
Purification of vaccine components and negative stain electron microscopy of nanoparticles. **Refers to Figure 1**. (A) Diagram of SpyCatcher003-mi3 NP construct. (B) SDS-PAGE with Coomassie staining of NP fractions after size exclusion chromatography purification (L-Ladder, F-Fraction). Gels converted to Coomassie blue color in PowerPoint. (C) SDS-PAGE with Coomassie staining of gel for products of SpySwitch purification of soluble gD (L-Ladder, SNT-Supernatant, FT-Flowthrough, W-Wash, E-Elution). Gels converted to Coomassie blue color in PowerPoint. (D) SDS-PAGE with Coomassie staining of gel for products of SpySwitch purification of soluble gH/gL (L-Ladder, SNT-Supernatant, FT-Flowthrough, W-Wash, E-Elution). Gels converted to Coomassie blue color in PowerPoint. (E) Negative stain electron microscopy micrographs of NP, gD NP, and gH/gL NP. Red boxes indicate insets shown in Figure 1D.

**Supplemental Figure 2.**
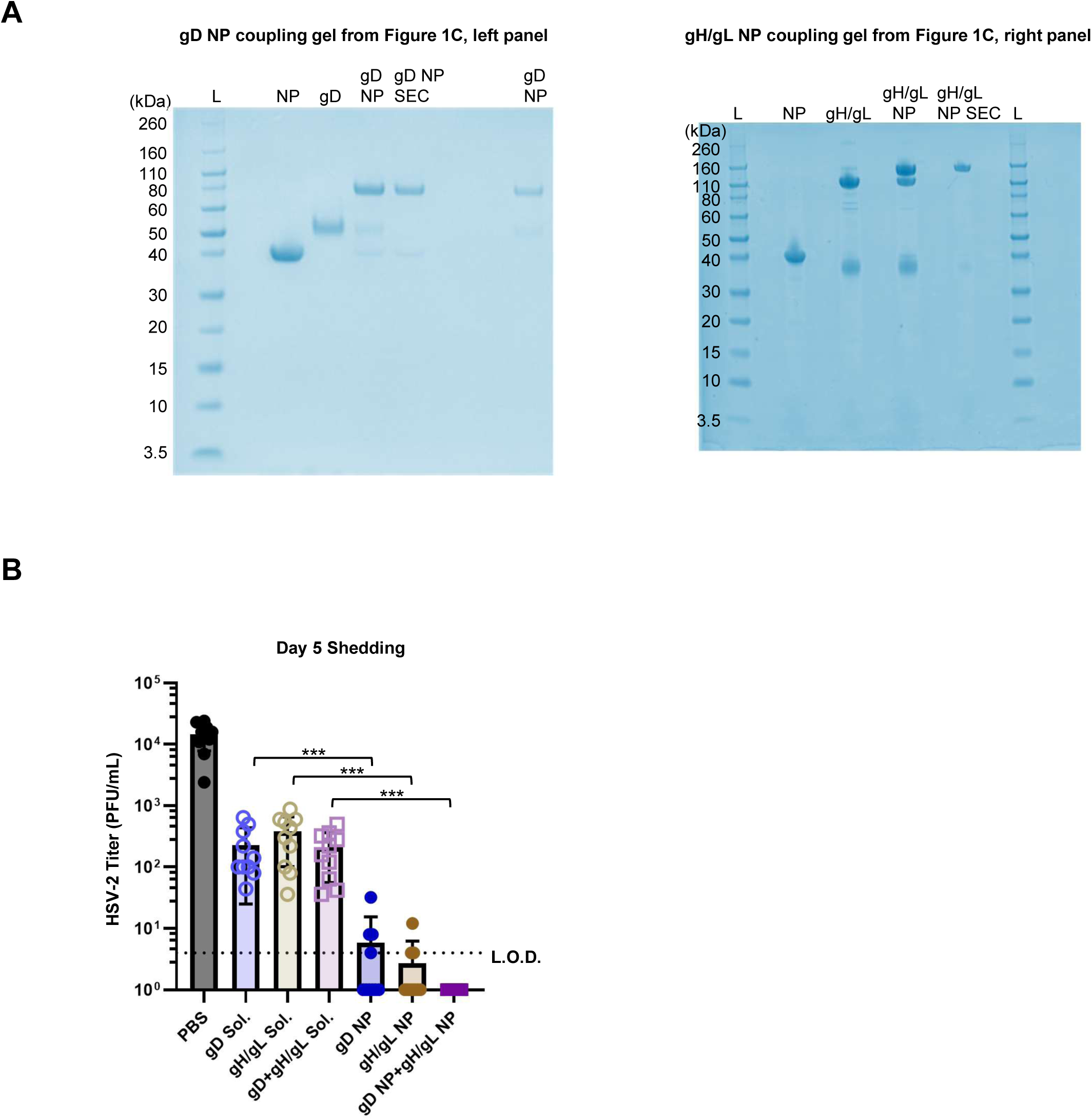
Full length gels presented in the paper. **Refers to Figures 1 and 2**. (A) Uncropped SDS-PAGE with Coomassie staining of antigen-NP coupling from Figure 1C (left panel is gD NP, right panel is gH/gL NP). Gels converted to Coomassie blue color in PowerPoint. (B) HSV-2 titers from vaginal swabs of individual mice on day 5 associated with Figure 2. Bars indicate means and error bars indicate standard deviations. L.O.D. is the limit of detection. *** for p<0.0001.

**Supplemental Figure 3.**
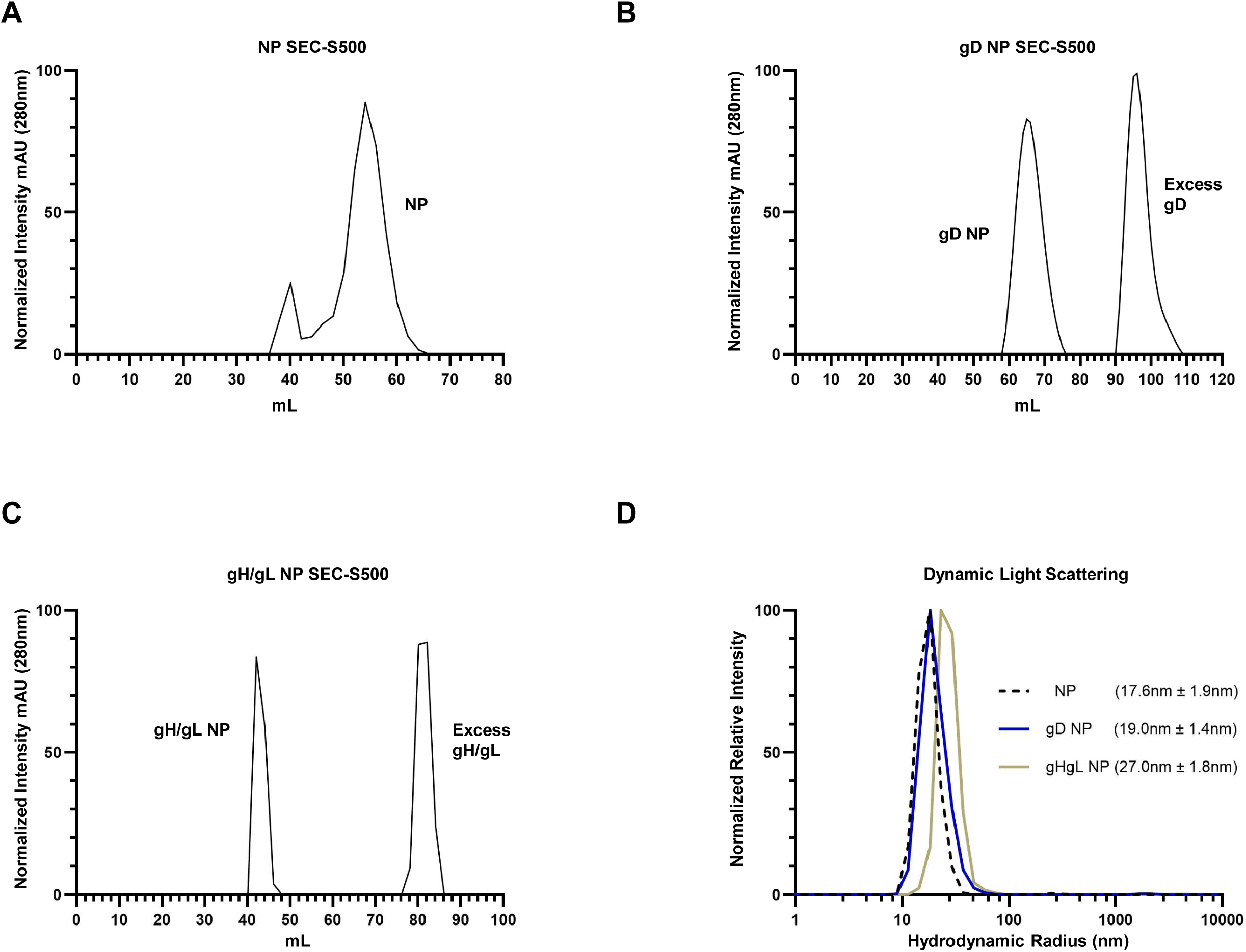
Purification of vaccine components and negative stain electron microscopy of nanoparticles. **Refers to Figure 1**. (A) Size exclusion chromatography (SEC) of NPs. (B) SEC of gD NP. The first peak is gD NP, and the second peak is the excess soluble gD. (C) SEC of gH/gL NP. The first peak is gH/gL NP, and the second peak is the excess soluble gH/gL. (D) Dynamic light scattering of uncoupled NP, gD NP, and gH/gL NP. Numbers represent hydrodynamic average radius with average polydispersion.

**Supplemental Figure 4.**
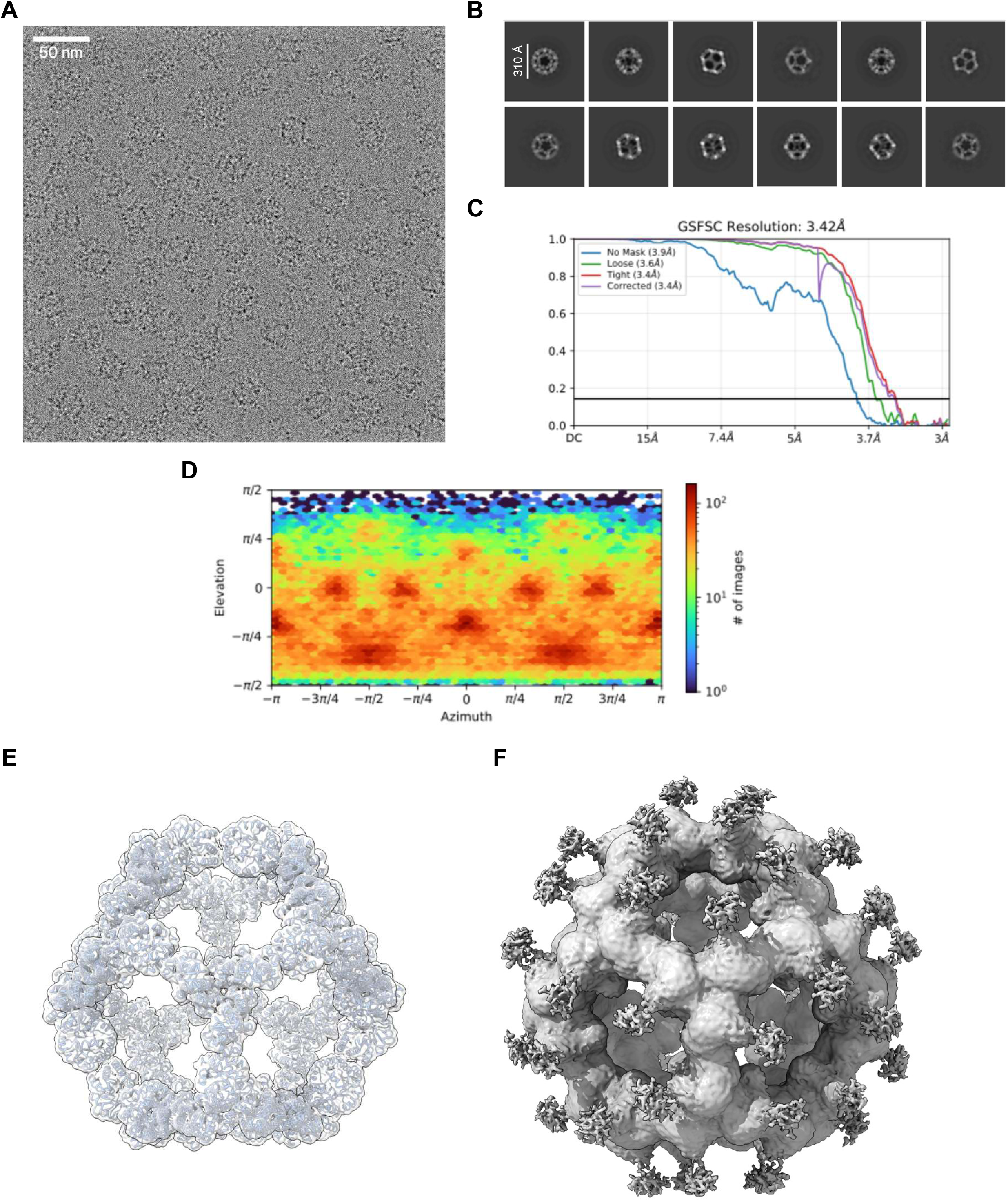
Cryogenic electron microscopy (CryoEM) of gH/gL NP. **Refers to Figure 1**. (A) Exemplary raw micrograph of gH/gL NPs. (B) Representative 2D classes of gH/gL NPs. (C) Fourier Shell Correlation (FSC) curves estimating the resolution of the reconstructed gH/gL NP 3D structure. (D) Viewing direction distribution plot of 3D structure. (E) CryoEM map (grey) at 10 σ level with PDB 7B3Y (blue) docked in the map. (F) CryoEM map (grey) at 0.8 σ level shows protrusions from NP subunits.

**Supplemental Figure 5.**
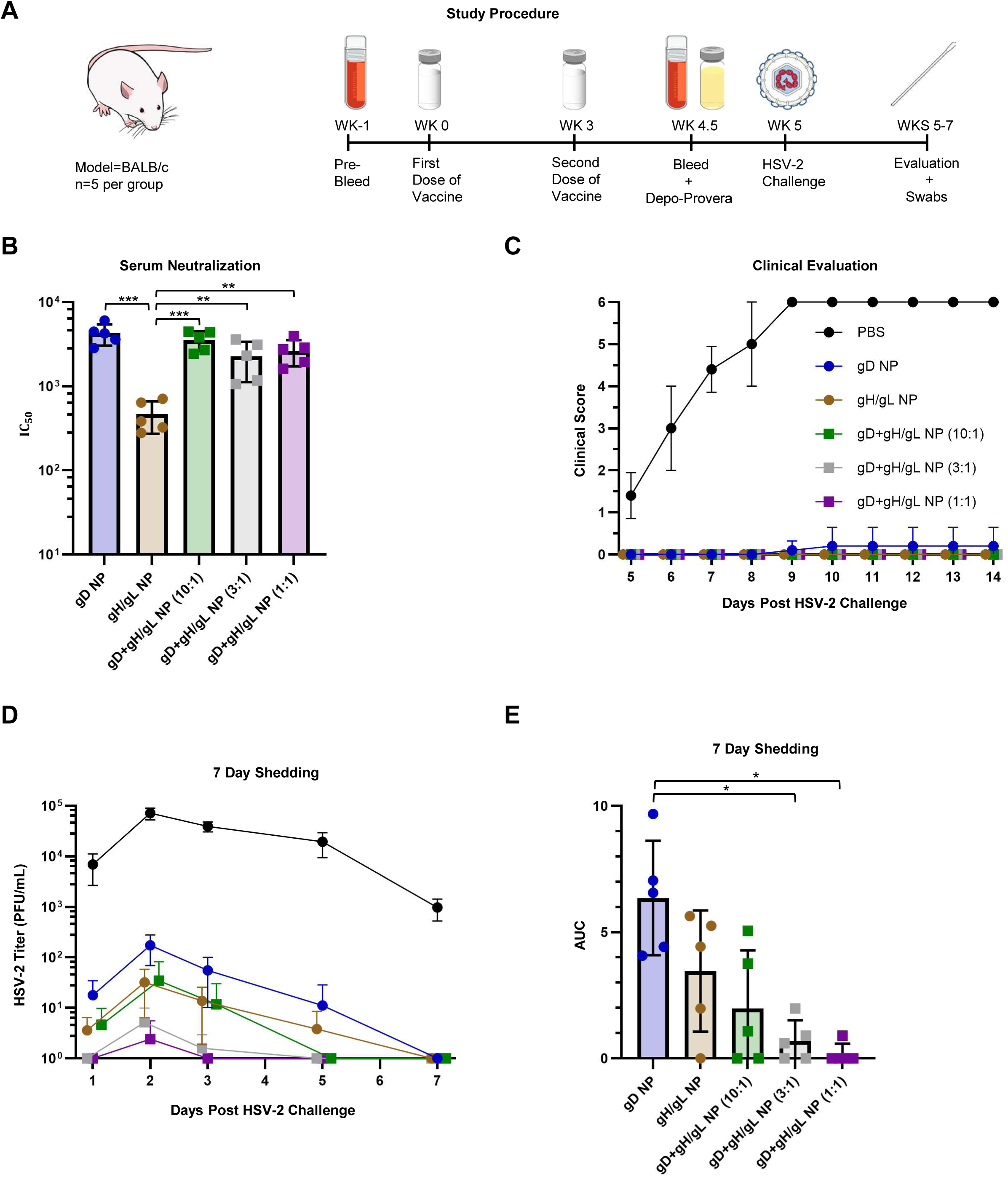
Protection of mice immunized with gD-NP, or gH/gL-NP, or combinations of gD-NP and gH/gL-NP at different ratios from HSV-2 vaginal challenge. **Refers to Figures 2 and 3**. BALB/c mice (n=5 per group) were immunized IM with 2 doses of constructs (5 µg) in SAS adjuvant, 3 weeks apart. Mice were intravaginally challenged with 64,000 PFU of HSV-2 strain 333. (A) Schematic of the study. Sera was analyzed from mice bled 1.5 weeks after the second dose of vaccine. (B) Neutralizing antibody titers in serum from vaccinated mice after dose 2. IC_50_ is the dilution of sera that inhibits infection by 50%. Bars indicate means and error bars indicate standard deviations. **p<0.0014, ***p<0.0001. (C) Mice were monitored daily for clinical signs of HSV-2 infection including vaginal erythema, genital lesions, genital hair loss, ruffled fur, lethargy, abnormal gait, hunched back, or hind-limb paralysis and given a score of 0 (no disease) to 6 (dead) (see Methods). Data points indicate means and error bars indicate standard deviations. There were no statistical differences between groups receiving NPs. (D) HSV-2 titers from vaginal swabs taken on days 1, 2, 3, 5, and 7 post-challenge. Data points indicate means and error bars indicate standard deviations. (E) Area under the curve (AUC) analysis of panel D showing 7 day shedding. Bars indicate means and error bars indicate standard deviations. *p<0.0071.

**Supplemental Figure 6.**
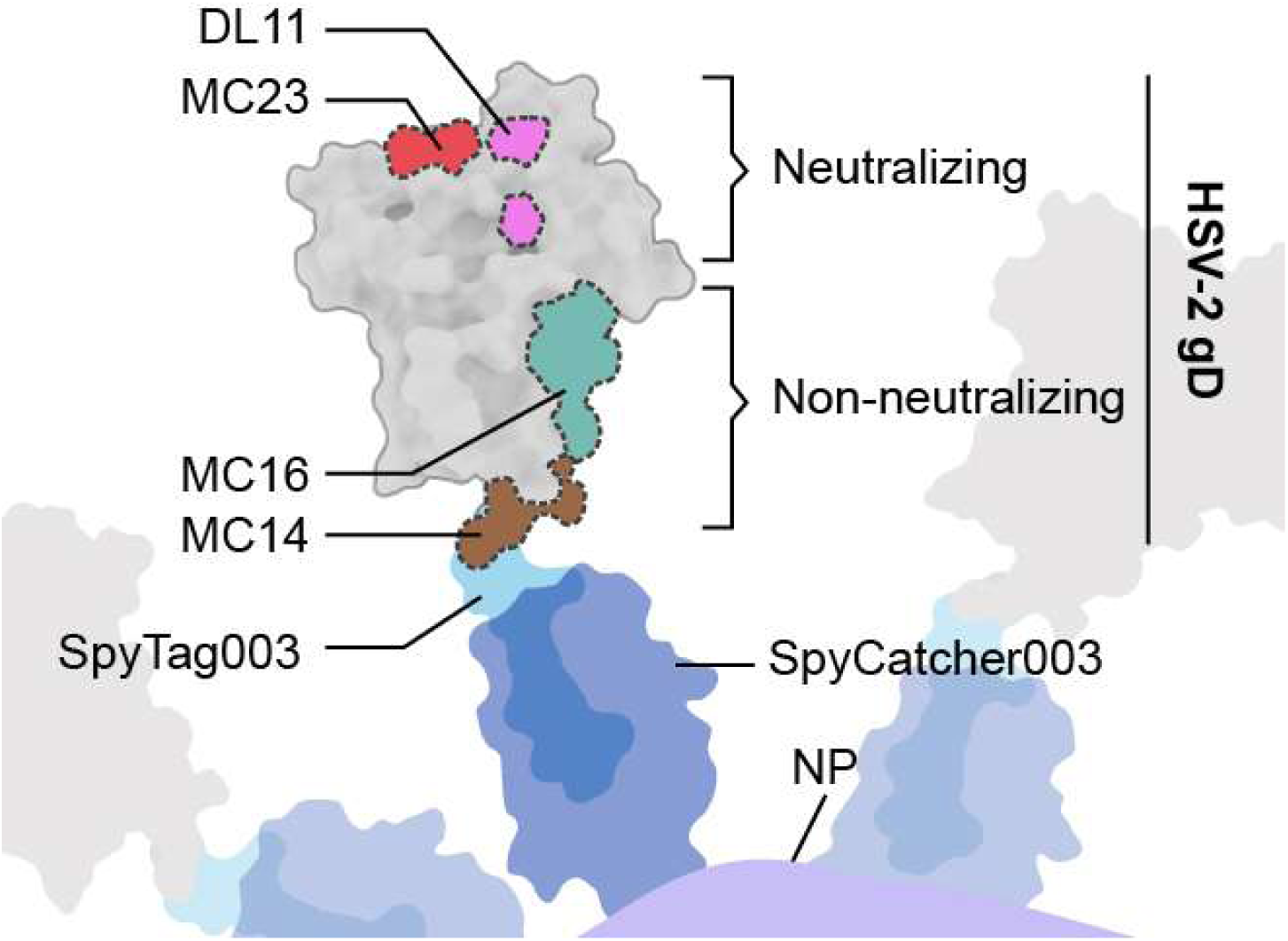
Schematic of immune focusing on receptor binding domains of HSV-2 gD enabled by NP display. **Refers to Figure 6**. The epitopes^27^ of monoclonal antibodies used for the sera competition assay in Figure 6 were mapped onto PDB (2C36) of HSV-1 gD (HSV-1 gD has more of its structure resolved than HSV-2 gD and shares significant amino acid homology). The carboxyl terminus of gD is less accessible since it is oriented towards the NP and epitopes near the amino terminus are freely exposed. This enforced orientation of gD may enable immune focusing on neutralizing epitopes and sterically hinder immune responses to non-neutralizing epitopes.

